# Mesostriatal dopamine is sensitive to specific cue-reward contingencies

**DOI:** 10.1101/2023.06.05.543690

**Authors:** Eric Garr, Yifeng Cheng, Huijeong Jeong, Sara Brooke, Laia Castell, Aneesh Bal, Robin Magnard, Vijay Mohan K. Namboodiri, Patricia H. Janak

## Abstract

Learning causal relationships relies on understanding how often one event precedes another. To gain an understanding of how dopamine neuron activity and neurotransmitter release change when a retrospective relationship is degraded for a specific pair of events, we used outcome-selective Pavlovian contingency degradation in rats. Two cues were paired with distinct food rewards, one of which was also delivered in the absence of either cue. Conditioned approach was attenuated for the cue-reward contingency that was degraded. Dopamine neuron activity in the midbrain and dopamine release in the ventral striatum showed a profile of changes in cue- and reward-evoked responding that was not easily explained by a standard reinforcement learning model. An alternative model based on learning causal relationships was better able to capture evoked dopamine responses during contingency degradation, as well as conditioned behavior following optogenetic manipulations of dopamine during noncontingent rewards. Our results suggest that mesostriatal dopamine encodes the contingencies between meaningful events during learning.

## Introduction

Temporal contiguity between events—how close they occur in time—is not sufficient to explain learning. In appetitive Pavlovian conditioning in which animals learn to anticipate rewards based on antecedent cues, learning can be stunted by free rewards in the absence of the cue (Delamater, 1995; Durlach & Shane, 1993; Ostlund & Balleine, 2007) even when contiguity between cues and rewards is held constant. A similar phenomenon holds for aversive Pavlovian conditioning, in which animals fail to freeze in response to a cue that predicts shock when shock is also delivered at the same rate in the absence of the cue (Rescorla, 1968). These findings have been used to argue that learning is a function of the cue-outcome contingency (Gallistal & Balsam, 2014).

Despite the importance that contingency has for learning, investigations of midbrain dopamine (DA) function emphasize a role in learning that is based on contiguity. A prominent theory of midbrain DA function comes from temporal difference reinforcement learning (TDRL), which uses prediction errors to update state and action values (Sutton & Barto, 2018). There is a remarkable resemblance between phasic DA activity and TDRL prediction errors (Amo et al., 2022; Kim et al., 2020; Schultz et al., 1997), and exogenously evoked DA modulates behavior as if evoking an artificial prediction error (Keiflin et al., 2019; Maes et al., 2020; Saunders et al., 2018; Steinberg et al., 2013). An assumption of TDRL is that agents learn values of cues and actions, where value is defined as the time-discounted expectation of future reward (Sutton & Barto, 2018). Put simply, the further away in time a reward is placed from some state, the less value will accrue to that state. Thus, value in TDRL is dependent on temporal contiguity between events, and many studies of how DA contributes to learning confound changes in value with changes in contingency. This is not to say that all variations of TDRL consider temporal contiguity as sufficient for learning (Sutton & Barto, 1990), but rather as a necessary condition.

In the current study, we sought to hold the contiguity between cues and rewards constant while degrading the contingency for one pair of events but not another. This is known as outcome-selective contingency degradation. Two distinct cues were followed by distinct rewards, but one reward was also delivered noncontingently during the inter-trial interval (ITI). From the point of view of a TDRL model that does not assign value to the ITI, the value of the degraded cue remains unchanged because it is followed by the same reward at the same interval and the same probability as the control cue. Even if the model assigns value to the ITI, under outcome-selective contingency degradation, the increased ITI value will affect both cues equally. TDRL thus predicts that outcome-selective contingency degradation should have no differential effects on degraded and nondegraded cues, both in terms of conditioned behavior or mesostriatal dopamine dynamics.

Here, we replicate the finding that outcome-selective contingency degradation gradually diminishes conditioned responding. Fiber photometry recordings of midbrain DA neuronal activity and of DA release in the ventral striatum reveal phasic responses to cues and rewards that diminish with contingency degradation, with the reward response losing its sensitivity to local reward history. Individual differences in performance are predicted by the phasic and tonic DA signal during the cue-outcome interval. We also show that non-contingent DA neuron activity during the ITI is sufficient, but not necessary, to attenuate conditioned responding using optogenetic approaches. Finally, we show that nearly all these results can be accounted for by a recently-described computational model where dopamine guides retrospective causal learning (Jeong et al., 2022).

## Results

### Contingency degradation attenuates conditioned responding for reward

To manipulate cue-reward contingency while holding contiguity constant, we used a procedure known as Pavlovian contingency degradation, wherein the contingency between a cue and a reward is diminished by delivering free rewards in the ITI. The design of the Pavlovian contingency degradation protocol was adapted from a previous report (Ostlund & Balleine, 2007). Rats learned two cue-reward associations during Pavlovian acquisition. Two auditory cues (tone and white noise) were presented in separate trials at random times and random order for 20 seconds followed by 0.5 probability of reward delivery (grain pellet or sucrose water). Each cue was associated with a different reward. During contingency degradation cues were still followed by probabilistic rewards, but one reward was delivered non-contingently during the ITI. This meant that the contingency between the reward delivered freely during the ITI and its associated cue was degraded, while the other cue-reward contingency remained intact. Put another way, one of the cues signaled a negligible change in reward rate (see description of cycle-to-trial ratios below).

The conditioned response was defined as the anticipatory head entry into the food port during the 20 second cue and before trial outcome. During the acquisition phase, the mean entry rate increased during both cues equally, but during contingency degradation, responding became weaker during the degraded cue compared to the non-degraded cue (**Figure 1B**). Repeated-measures ANOVAs on conditioned entry rates with and without baseline entries subtracted revealed two-way interactions between cue type and training phase (*F*’s(1,34) > 18.24, *p*’s < .001) and three-way interactions between cue type, session, and training phase (*F*’s(7,238) > 2.64, *p*’s < .013). Post-hoc contrasts revealed a difference between cue types during the last four combined sessions of contingency degradation (*p* < .05). There was also a main effect of training phase whether baseline was subtracted or not (*F*’s(1,34) > 4.99, *p*’s < .033), indicating that, in addition to cue-selective effects, contingency degradation also generally weakened conditioned responding. The effect of contingency degradation was not mediated by context, which rules out a value-based account of contingency degradation whereby the value of the context overshadows the value of the degraded cue (**Supplemental Figure 1D**).

**Figure 1.**
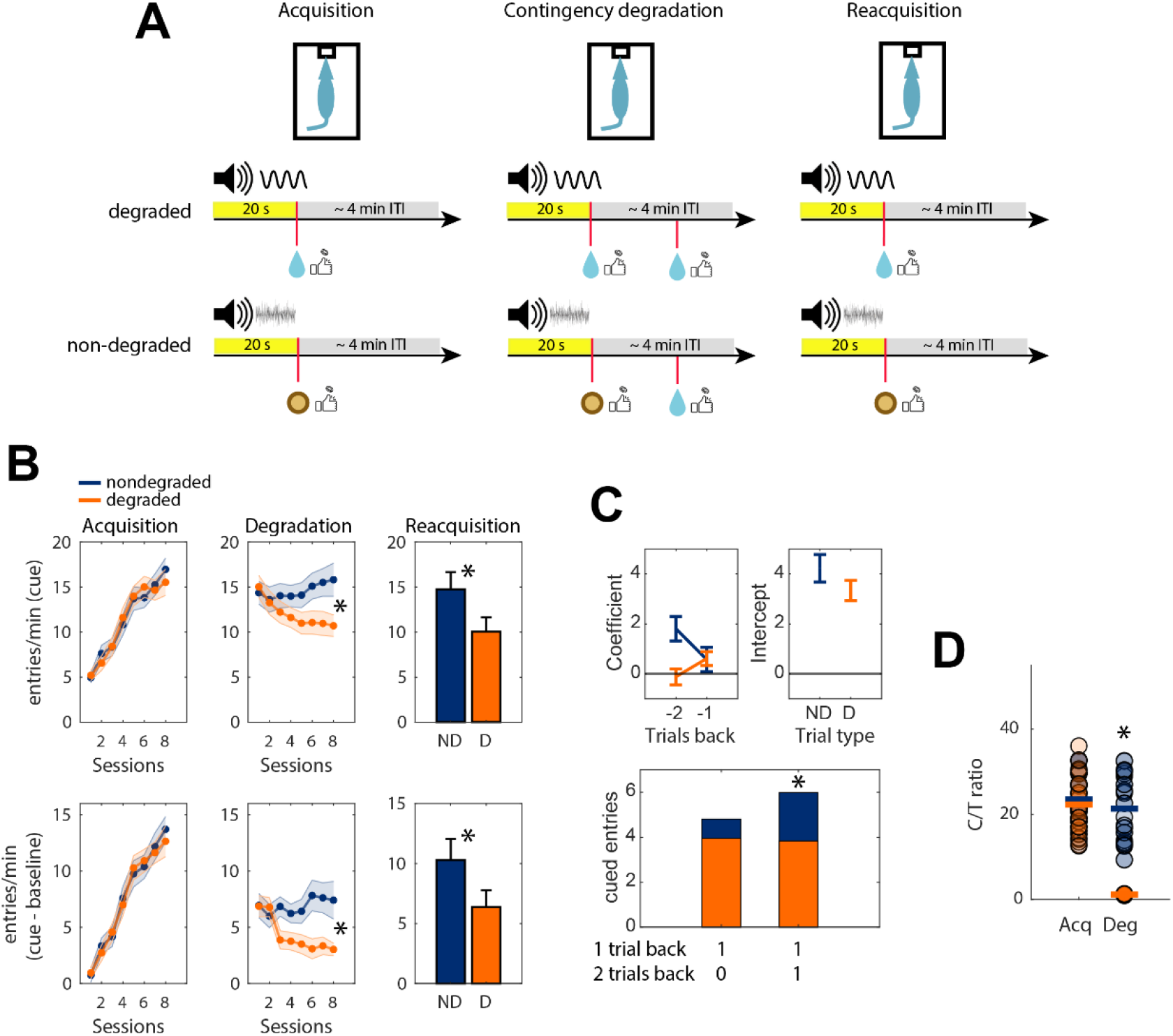
Conditioned responding during Pavlovian contingency degradation. **(A)** Design of Pavlovian contingency degradation experiments showing the two trial types with different food rewards. Trials were presented in random order. The specific cue-outcome and non-contingent outcome assignments are shown for illustration purposes, but they were counterbalanced across rats. Coin flip icon means 0.5 probability of reward. **(B)** Mean port entry rates during the 20 second cue period (top) and with baseline subtracted (bottom) for each phase of the experiment. Shaded boundaries are standard errors of the mean. Grey lines are individual rats. Data are pooled across 35 rats from 3 separate experiments (VTA GCaMP, NAc dLight, and context change/reward preference), except during reacquisition, which only includes rats from the VTA GCaMP and NAc dLight experiments. **(C)** Top: mean regression coefficients (left) and intercepts (right) when the number of cued port entries was regressed against trial outcome history, shown separately for nondegraded and degraded trials. Bottom: mean predicted entry rates on trial *n* as a function of the outcome on trials *n*-1 and t *n*-2. 1 indicates reward and 0 indicates omission. All data are taken from the last session of contingency degradation. **(D)** Cycle-to-trial ratios calculated from the final sessions of acquisition and contingency degradation. Circles are individual rats and horizontal bars are means. Data are pooled across 27 rats from the VTA GCaMP and NAc dLight experiments.

To confirm that contingency degradation affected learning and not just performance, a reacquisition session was run in which the original contingencies from the acquisition phase were reinstated (**Figure 1B**). The mean entry rate was significantly higher during the non-degraded cue whether baseline was subtracted (*t*(26) = 2.17, *p* = .04) or not (*t*(26) = 2.58, *p* = .016).

Next, we examined how local trial history affected conditioned port entries. This was achieved by modeling the number of cued port entries as a linear combination of trial outcomes up to two trials back in time. The regression coefficients and intercepts were combined for each rat to yield the predicted number of cued entries as a function of trial history (**Figure 1C**). A repeated-measures ANOVA revealed a main effect of reward history (*F*(1,34) = 4.23, *p* = .047), a main effect of cue type (*F*(1,34) = 5.57, *p* = .024), and a cue type x reward history interaction (*F*(1,34) = 4.11, *p* = .05). Post-hoc contrasts showed that two consecutively rewarded trials increased the number of port entries more for the subsequent nondegraded cue (*p* < .05). However, when a rewarded trial was preceded by an omission trial, the subsequent cue port entries did not significantly differ between cue type (*p* > .05). This analysis shows that behavioral sensitivity to local reward history was weaker for the cue that underwent contingency degradation.

To confirm that the non-degraded cue signaled a larger change in reward rate, the cycle-to-trial (C/T) ratio was computed (**Figure 1D**). The ‘cycle’ is the mean interval between all deliveries of one reward type, and the ‘trial’ is the mean interval between that same reward and its preceding cue during a trial (Balsam et al., 2010). During contingency degradation, the C/T ratio was significantly smaller for the degraded versus non-degraded cue (*t*(26) = 13.90, *p* < .001). These analyses show that conditioned head entry rates during a Pavlovian cue were selectively attenuated by outcome-selective contingency degradation.

### Contingency degradation attenuates the VTA DA neuron response to cue and reward

To determine the impact of contingency degradation on encoding of cue-reward contingencies, we measured DA neuron activity in the ventral tegmental area (VTA) when one cue-reward contingency was degraded while the other remained intact. Tyrosine hydroxylase (TH)-Cre rats (*n* = 13) were injected with a Cre-dependent GCaMP6f virus in the VTA for imaging of calcium fluorescence via fiber photometry. The ΔF/F signal was aligned to cue onset, reward delivery, and reward omission during the last session of contingency degradation (**Figure 2B**). The DA response to the non-degraded cue was significantly higher than to the degraded cue (*t*(12) = 2.36, *p* = .037), and the DA response to reward onset was greater following the non-degraded cue than the degraded cue (*t*(12) = 3.82, *p* = .002). The DA response to reward omission was divided into a positive phase, when the signal rose above baseline, and a negative phase when the signal dropped below baseline (see Methods). The omission response did not differ in either the positive (*t*(12) = 2.13, *p* = .055) or negative phase (*t*(12) = 0.84, *p* = .42) when comparing the degraded and nondegraded outcome pair. During the first session of contingency degradation, the reward responses differed in the same direction but the cue responses did not, indicating that it is the cue-evoked component that changes with conditioned responding (**Supplemental Figure 2**). These findings show that DA neuron responses to cues and rewards diminish during contingency degradation.

**Figure 2.**
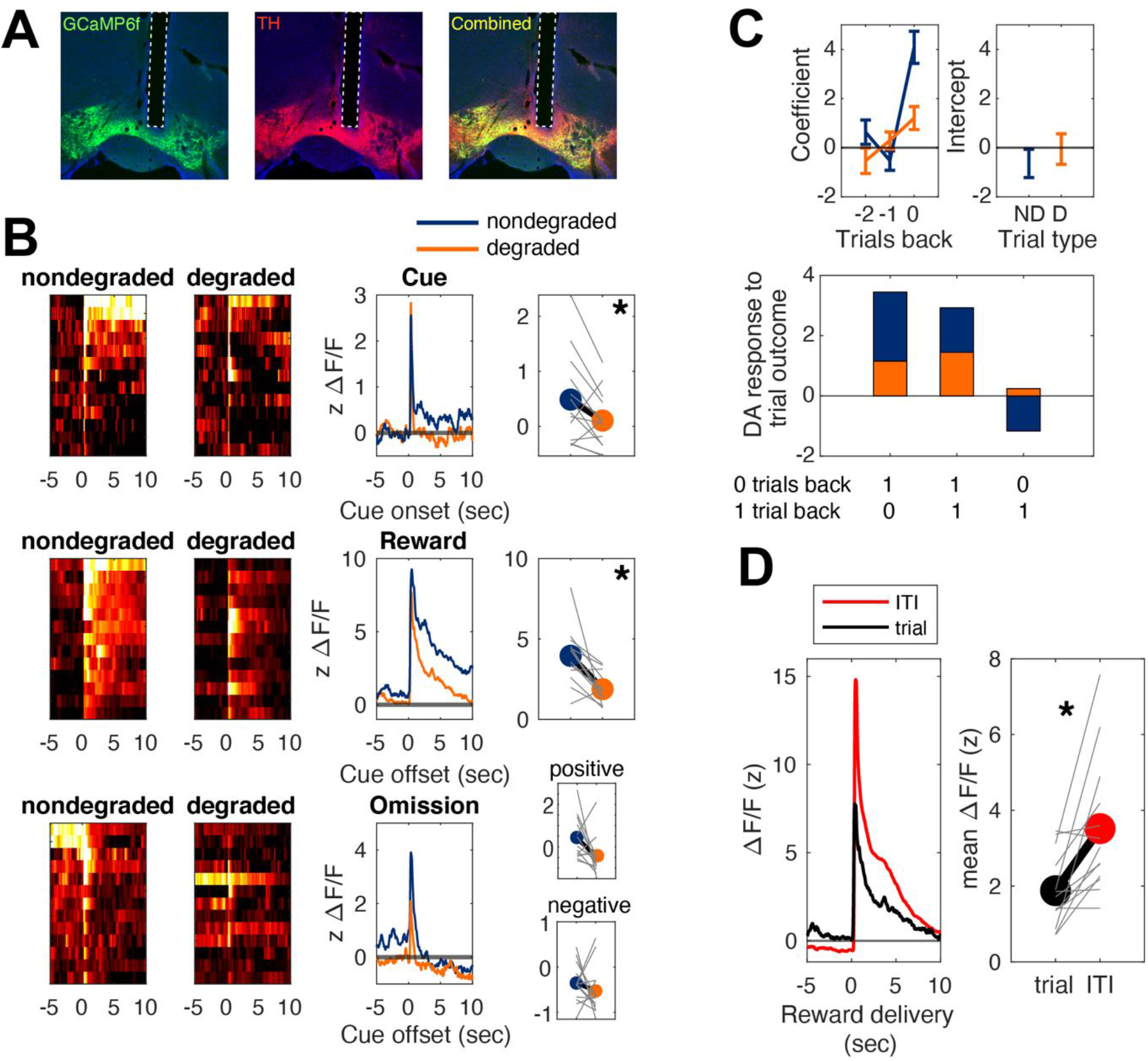
Fiber photometry recordings in VTA DA neurons during contingency degradation. **(A)** Coronal section from one TH Cre rat injected with DIO-GCaMP6f in the VTA. Dotted lines outline the fiber tract. **(B)** Left column: heat maps showing GCaMP response aligned to trial events separated by nondegraded and degraded trials. Each row shows the mean for each rat. Middle column: Mean photometry traces in response to trial events, shown separately for nondegraded (blue) and degraded (orange) trials. Right column: mean z-scored ΔF/F in response to trial events for nondegraded and degraded trials. Reward omission means were quantified separately for positive (top) and negative (bottom) phases of the signal. Grey lines show individual rats. All data are from the last session of contingency degradation. **(C)** Top: mean regression coefficients (left) and intercepts (right) when the GCaMP response to trial outcome was regressed against trial outcome history, shown separately for nondegraded and degraded trials. Bottom: mean predicted GCaMP response at the time of the outcome on trial *n* as a function of the outcome on trial *n* and trial *n*-1. 1 indicates reward and 0 indicates omission. All data are taken from the last session of contingency degradation. **(D)** Mean GCaMP signal aligned to the same reward delivered during the ITI or at the end of a trial at the offset of the degraded cue. Grey lines represent individual rats. All data are from the last session of contingency degradation.

We ran additional analyses to explore whether the observed DA signal conformed to a canonical reward prediction error signal. One way to test for reward prediction error encoding is to quantify how the neural response to trial outcome is affected by local reward history (Bayer & Glimcher, 2005; Ottenheimer et al., 2020; Parker et al., 2016). This was achieved by modeling the photometry response to the trial outcome as a linear combination of trial outcomes starting from any given trial and going two trials back. The regression coefficients and intercepts were combined for each rat to yield the predicted photometry response as a function of trial history (**Figure 2C**). A repeated-measures ANOVA revealed a main effect of reward history (*F*(2,24) = 40.83, *p* < .001), no main effect of cue type, and a cue type x reward history interaction (*F*(2,24) = 15.48, *p* < .001). Post-hoc contrasts showed that the outcome response following the non-degraded cue resembled the pattern that would be expected if DA neurons were computing prediction errors. Specifically, the photometry response was greatest when the current trial was rewarded but the previous trial was not, followed by a smaller response when reward was delivered two trials in a row, and an even smaller response when the current trial was not rewarded by the previous one was (*p* < .05). This was not true of the outcome response following degraded trials (*p* > .05). The same analysis was applied to the photometry response to cue onset, with no main effects or interaction (**Supplemental Figure 4A**).

A reward prediction error account of DA signaling predicts that the reward delivered during the ITI should elicit a larger response compared to when it is delivered after a fixed-duration cue. We compared the VTA DA response aligned to reward delivery at the end of a trial—specifically, at the offset of the degraded cue—and to delivery during the ITI (**Figure 2D**). Notably, the reward identity is the same in both cases. The response to the reward delivered during the ITI was larger than to the same reward delivered at the end of a trial (*t*(12) = 3.56, *p* = .004).

### Contingency degradation attenuates NAc DA release to cue, reward delivery, and reward omission

To confirm that these findings hold for DA neurotransmitter release, wildtype rats (*n* = 14) were injected with a dLight1.2 virus in the core of the nucleus accumbens (NAc) for imaging of dopamine release via fiber photometry. The z-scored ΔF/F signal aligned to trial events is shown in **Figure 3B**. Similar to VTA DA neuron activity, DA release aligned to the non-degraded cue was significantly higher than to the degraded cue (*t*(13) = 3.18, *p* = .007), and DA release aligned to reward onset was greater following the non-degraded cue than the degraded cue (*t*(13) = 5.54, *p* < .001). The reward omission response did not differ during the positive phase (*t*(13) = 1.67, *p* = .118), but was significantly different during the negative phase (*t*(13) = 2.45, *p* = .029), with a greater drop below baseline after termination of the non-degraded cue. During the first session of contingency degradation, the reward responses differed in the same direction but the cue responses did not (**Supplemental Figure 3).**

**Figure 3.**
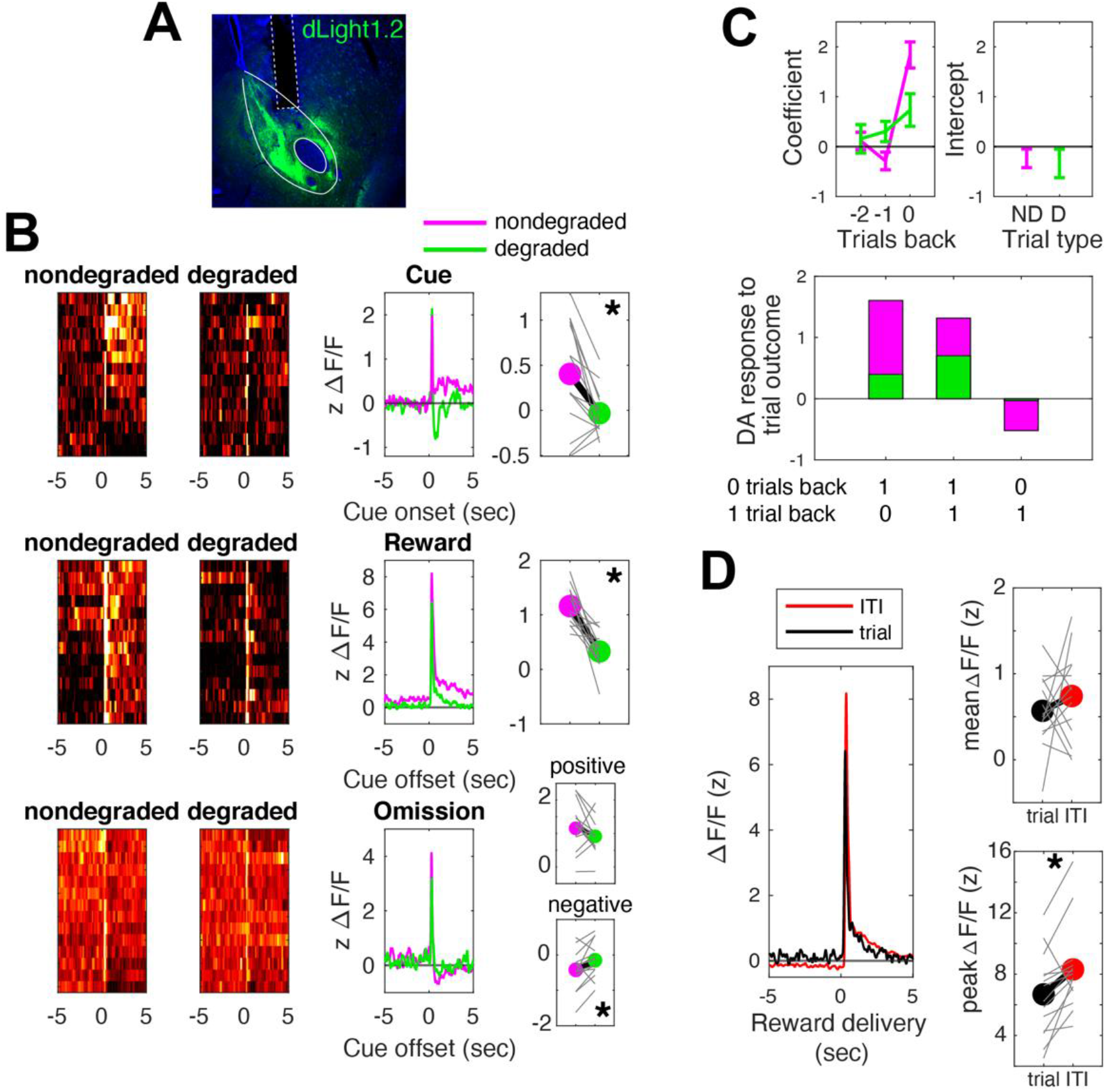
Fiber photometry recordings of DA release in NAc during contingency degradation. **(A)** Coronal section from one rat injected with dLight1.2 in the NAc core. Dotted lines outline the fiber tract. **(B)** Same as Figure 2, except nondegraded trials are in magenta and degraded trials are in green. **(C)** Same as in Figure 2. **(D)** Same as in Figure 2, but also showing the peak z ΔF/F signal (right). All data are from the last session of contingency degradation.

As above, DA release during trial outcome was modeled as a combination of current and past trial outcomes (**Figure 3C**). The predicted response to trial outcomes as a function of trial history showed a main effect of reward history (*F*(2,26) = 15.31, *p* < .001), a main effect of cue type (*F*(1,13) = 9.48, *p* = .009), and a cue type x reward history interaction (*F*(2,26) = 7.55, *p* = .003). Post-hoc contrasts showed that the outcome response following the non-degraded cue was greatest when the current trial was rewarded but the previous trial was not, followed by a smaller response when reward was delivered two trials in a row, and an even smaller response when the current trial was not rewarded but the previous one was (*p* < .05). This was not true of the outcome response following degraded trials (*p* > .05). This analysis shows that the modulation of NAc DA by trial outcomes accords with TDRL only when the outcome is contingent on the cue, as observed above for the VTA GCaMP analysis.

We next compared DA release aligned to reward delivery at the end of a trial and to delivery during the ITI (**Figure 3D**). The mean response to the reward delivered during the ITI did not differ from the response to the same reward delivered at the end of a trial (*t*(13) = 0.34, *p* = .743), but the peak response did (*t*(13) = 2.96, *p* = .011). Overall, the measurements of DA cell body activity and DA terminal release reveal a highly similar pattern of changes in cue- and reward-elicited activity following contingency degradation that are specific to the targeted cue-outcome pair. These findings confirm that DA signaling is sensitive to changes in contingency.

### Neither behavior nor DA transients are explained by satiety

The differences in the behavioral and photometry responses to events during degraded and non-degraded cue presentations could be explained by differences in satiety. Specifically, during contingency degradation one reward type is delivered more frequently than the other, and it is possible that rats could become sated on that reward and this could differentially affect both the behavioral response and the DA response to trial events. Reward preference tests showed that contingency degradation induced only general satiety, not specific satiety (**Supplemental Figure 1E**). To test whether the photometry response to trial events was affected by satiety, we regressed the photometry response to cues, reward deliveries, and reward omissions on the times at which those events occurred during the last degradation session. The assumption is that satiety should grow over the course of the session. The times at which cues, reward deliveries, and reward omission occurred did not affect the photometry responses to those events for either VTA DA neuron or NAc DA release recordings (**Supplemental Figure 5**; one-sampled *t*-test *p*’s > .05).

An additional analysis of the DA reward response as a function of trial (first and last) and cue type (nondegraded and degraded) revealed only a main effect of cue type (*p*’s < .003), but no main effect of trial number nor an interaction. These results show that DA signals do not progressively decrease over session time as might be expected if satiety were contributing to the observed DA responses.

### Phasic and tonic DA during the cue-outcome interval predicts conditioned responding

We observed that degrading the cue-reward relationship by adding noncontingent reward deliveries altered DA responses to both cue and reward and altered behavioral responding to the degraded cue. To relate the neural dynamics to behavioral performance, we first regressed the rate and timing of conditioned entries against the photometry responses to cues, rewards, and omission on a trial-by-trial basis. These analyses did not yield any meaningful patterns (**Supplemental Figures 6 and 7**). We then asked if there was a reliable relationship between the relative behavioral response to the two cues and the neural responses to the two cues. Rats were split into two groups: those that showed a behavioral effect of contingency degradation, and those that did not. Groups were determined by subtracting the mean entry rate during the degraded cue from the mean rate during the non-degraded cue and using the 0 boundary to divide the groups. Given that a minority of rats did not show an effect of contingency degradation, analyses were combined across GCaMP and dLight experiments (see **Figure 4A and 4B** for individual experiment data). The cue-evoked response was segmented into phasic and tonic components (see Methods). These components were both predictive of behavioral performance (see **Supplemental Figure 8** for analysis of reward and omission responses, which were not predictive of performance).

**Figure 4.**
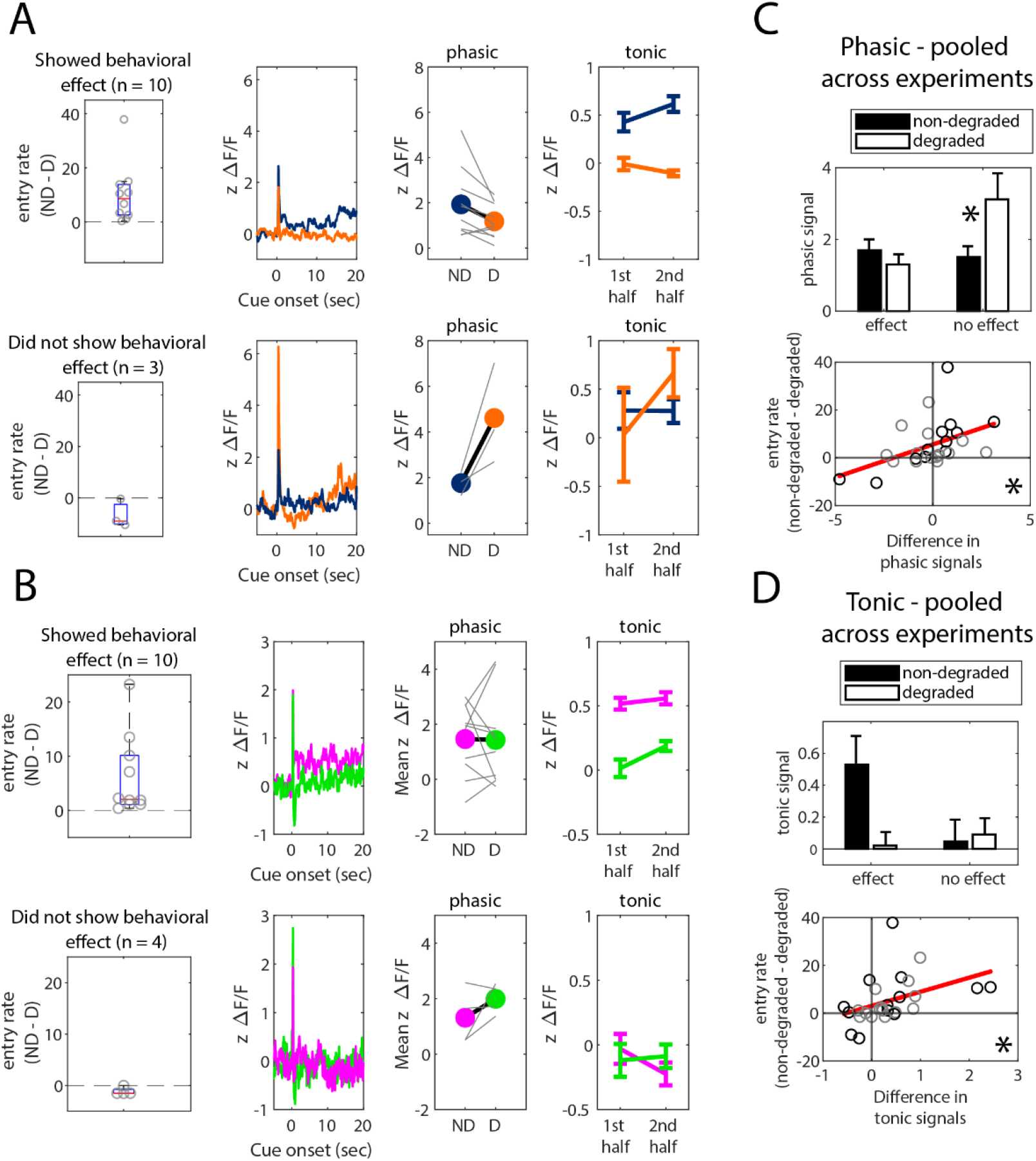
Phasic and tonic components of cue-evoked DA activity and release predict conditioned responding during contingency degradation. **(A)** Data from the VTA GCaMP experiment broken down by rats that did and did not show behavioral sensitivity to contingency degradation. Photometry data only from the cue-outcome interval is shown. **(B)** Same as panel A, but for the NAc dLight experiment. **(C)** Top: The phasic response to cue onset was analyzed as a function of cue type and group. Bottom: The difference in the phasic responses to degraded and non-degraded cues was used to predict behavior on a continuous measure. Black points come from the VTA GCaMP experiment and grey points are from the NAc dLight experiment. **(D)** Same as panel C, but for tonic DA during. All data are from the last session of contingency degradation.

For the phasic component of the DA response, a cue x group ANOVA yielded a significant interaction (**Figure 4C, top**; *F*(1,25) = 10.53, *p* = .003), and regressing the session-averaged difference in the entry rates between non-degraded and degraded trials against the difference between the phasic signals revealed a significantly positive relationship (**Figure 4C, bottom**; β = 2.76, *t*(25) = 2.61, *p* = .015). For the tonic component of the DA response, a cue x group ANOVA yielded an interaction that was shy of significance (**Figure 4D, top**; *F*(1,25) = 3.52, *p* = .072), but regressing the difference in the entry rates between non-degraded and degraded trials against the difference between tonic signals revealed a significantly positive relationship (**Figure 4D, bottom**; β = 5.90, *t*(25) = 2.33, *p* = .028). Overall, these data indicate that the phasic and tonic DA signals during the cue-outcome interval are sensitive to cue-reward contingency, and may drive individual differences in Pavlovian learning and motivation.

### Inhibiting DA activity and release during ITI rewards does not block the effect of contingency degradation on conditioned responding

During photometry recordings, non-contingent food rewards evoked a phasic response in VTA DA neurons and DA release in the NAc. To test whether non-contingent DA activity is necessary for rats to learn about contingency degradation, TH-Cre rats (*n* = 10) underwent optogenetic inhibition of DA neurons during non-contingent reward deliveries. Green laser stimulation was targeted to eNpHR-expressing DA neurons in the VTA only during non-contingent rewards during the ITI. A control group of WT rats (*n* = 9) also received the same conditioning and laser treatments. Conditioned entry rates across acquisition and degradation are shown in **Figure 5C**. Rats in both groups showed lower conditioned responding during the degraded vs. nondegraded cue (training phase x cue interaction (*F*(1,17) = 9.19, *p* = .008), with posthoc contrasts showing lower entry rates during the degraded versus non-degraded cue during contingency degradation (*p* < .05) but not acquisition (*p* > .05). However, there was no training phase x cue x group interaction (*F*(1,17) = 0.06, *p* = .808), suggesting that the optogenetic manipulation had no effect on conditioned entries.

**Figure 5.**
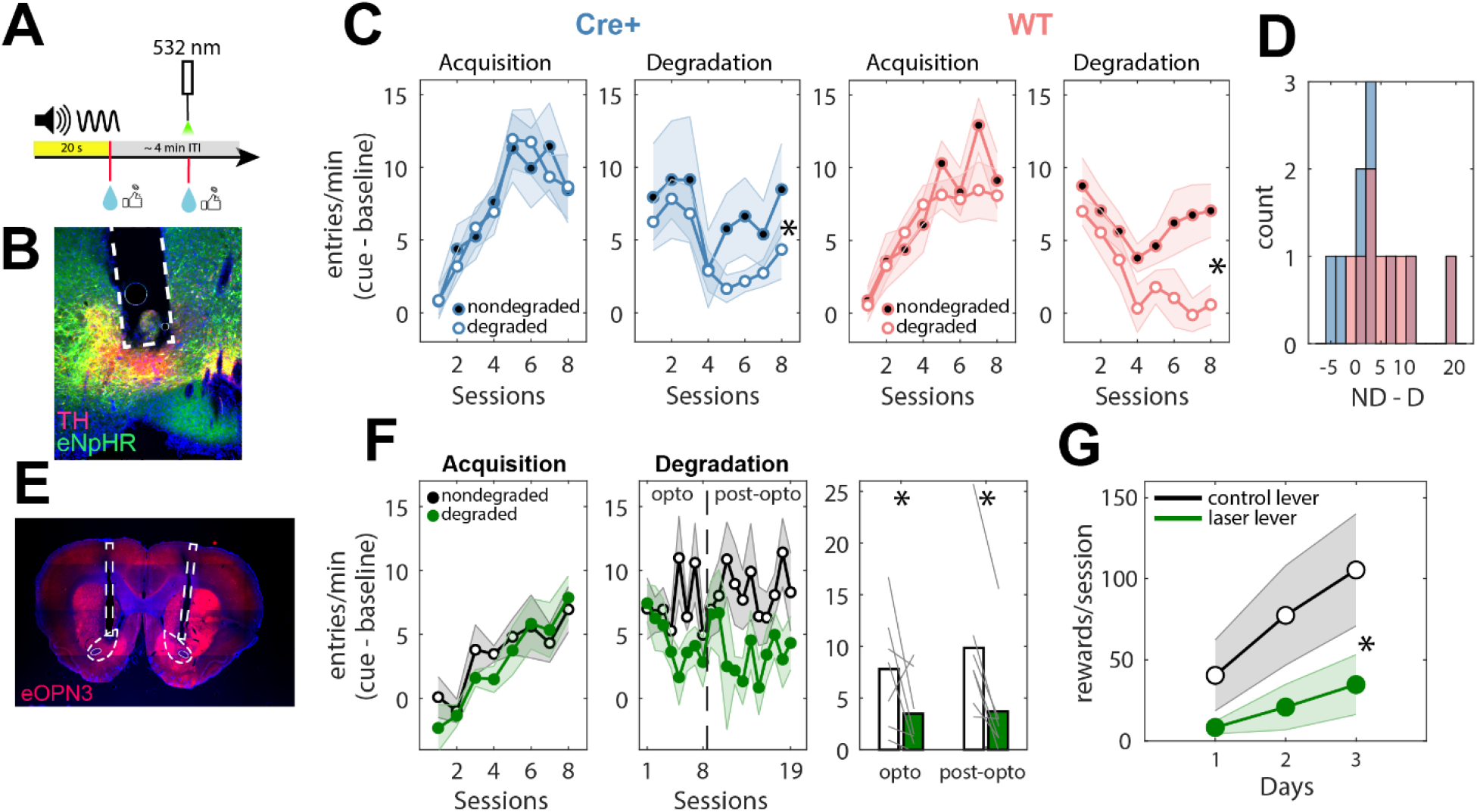
Using optogenetic inhibition to test whether non-contingent dopamine is necessary to learn about contingency degradation. **(A)** Green laser onset occurred during the presentation of only non-contingent rewards during the ITI, but not contingent rewards at trial offset. The illustration shows only the degraded trial type, but non-degraded trials were also intermixed. **(B)** Coronal section from one TH-Cre rat injected with DIO-eNpHR in the VTA and implanted with bilateral fibers in the VTA. Blue = DAPI, red = TH, yellow = eNpHR. **(C)** Port entry rates during each cue, with baseline subtracted. Shaded boundaries are standard errors of the mean. Steel blue = Cre+, light coral = WT. **(D)** Distribution of the differences between entry rates during nondegraded (ND) and degraded (D) cues, shown for each group. **(E)** Coronal section from one TH-Cre rat injected with SIO-eOPN3 in the VTA and implanted with bilateral fibers in the NAc core. Blue = DAPI, red = eOPN3 terminals. **(F)** Port entry rates during the cue for each phase of the experiment. The vertical dashed line in the middle figures represents the point at which laser stimulation was removed from the experimental sessions. Grey lines in the right-most graph are individual rats. **(D)** Acquisition of lever pressing on an FR1 schedule for rats expressing eOPN3. Both levers were associated with pellet rewards, but only one was associated with green laser stimulation.

We also optogenetically inhibited DA release in the NAc during non-contingent rewards. VTA DA axons in the NAc expressing the synaptic terminal-inhibiting opsin, eOPN3, were targeted with blue light during noncontingent reward deliveries during the ITI (see *Methods*). This experiment did not include a WT control group, and instead Cre+ rats underwent 8 sessions of contingency degradation with optogenetic inhibition (‘opto’) followed by 11 sessions without inhibition (‘post-opto’). Like the result of the previous experiment, optogenetic inhibition during non-contingent rewards did not affect entry rates (**Figure 5F**; main effect of cue: *F*(1,6) = 18.34, *p* = .005; no cue x training phase interaction: *F*(1,6) = .42, *p* = .540).

Unlike eNpHR, for which there has been extensive documented use in VTA DA neurons (Chang et al., 2016; Luo et al., 2018; McCutcheon et al., 2014; Sharpe et al., 2017), eOPN3 is a relatively new opsin (Mahn et al., 2021). To confirm the functionality of eOPN3, rats were next trained to press levers for pellet rewards. One lever delivered pellets and occasionally triggered laser stimulation (see *Methods*), while the other lever only delivered pellets. Rats showed a deficit in learning to press the laser-paired lever (**Figure 5G**; main effect of lever: *F*(1,6) = 7.09, *p* = .037). This result suggests that the inhibitory opsin was functional.

### Non-contingent VTA DA neuron activation attenuates a component of conditioned responding

The previous experiment shows that non-contingent DA neuron activity is not necessary to attenuate conditioned responding during contingency degradation. We next asked whether DA neuron activity is sufficient to attenuate conditioned responding during a variation of contingency degradation, using DA neuron stimulation as the trial outcome. TH-Cre+ rats were first trained to associate a 7-sec compound cue with unilateral VTA DA neuron optogenetic stimulation (1 sec, 20 Hz, delivered at cue offset) for twelve daily sessions (**Figure 6B**). TH-Cre-control rats that did not express ChR2 were run in parallel. We previously reported that pairing localizable cues with unilateral VTA DA neuron stimulation results in conditioned locomotion in rats—specifically, approach toward the cue and full body rotation contralateral to the stimulated hemisphere (Saunders et al., 2018). Therefore, we used DeepLabCut (Mathis et al., 2018) to track rat body parts during this procedure to detect any possible changes in behavior during cue-DA stimulation pairings and contingency degradation.

**Figure 6.**
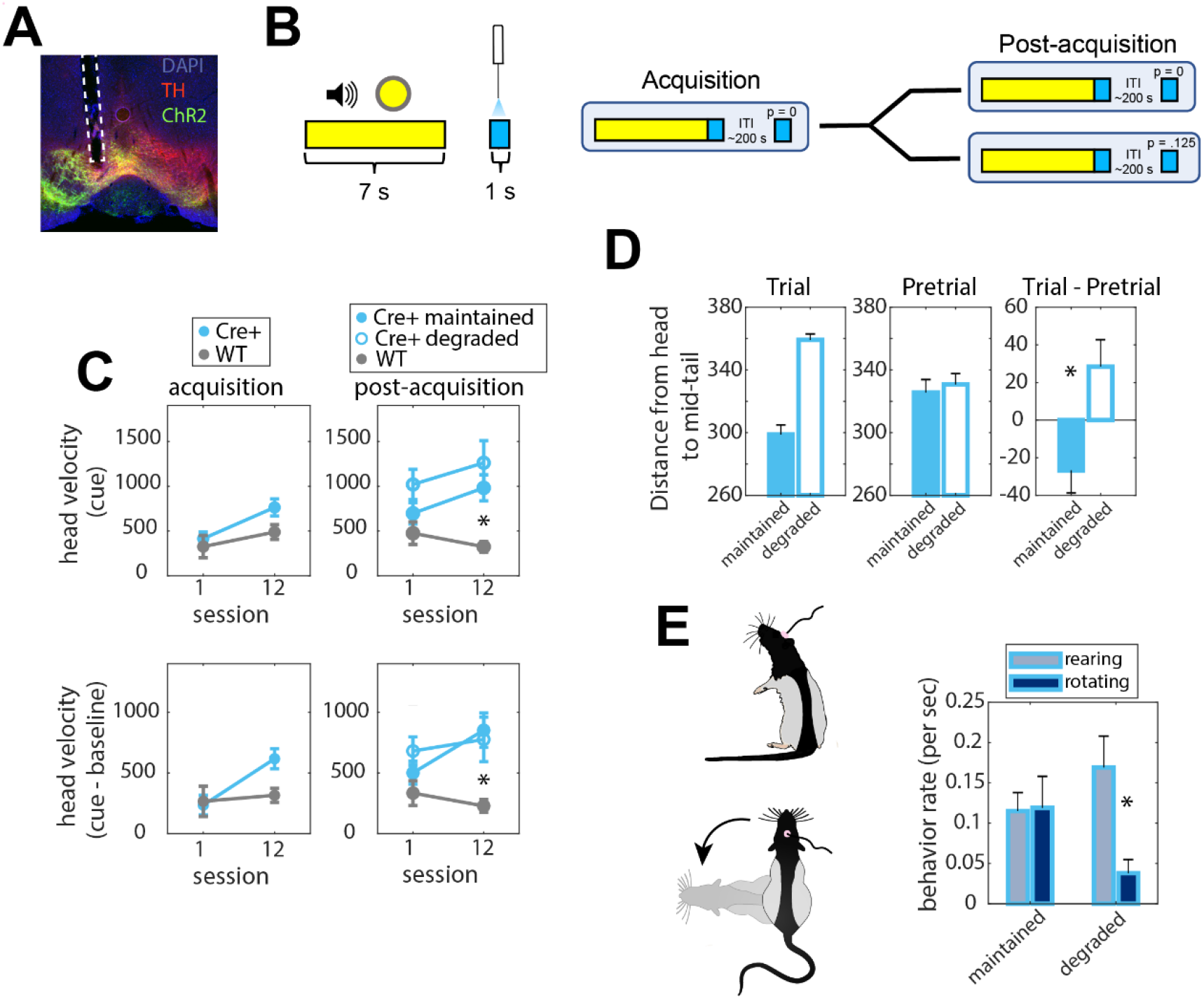
Using optogenetic stimulation to manipulate the contingency between a cue and VTA DA neuron activity. **(A)** Coronal section from one TH Cre rat injected with DIO-ChR2 in the VTA. Dotted lines outline the fiber tract. **(B)** Experiment design. During post-acquisition, some rats were maintained on the same conditioning protocol (top) or switched to contingency degradation (bottom). **(C)** Group velocity means during the cue period (top) and with baseline subtracted (bottom). **(D)** The group mean distances, in pixels, between the head and the middle of the tail during the trial (left), before the trial (middle), and as a difference between the trial and pretrial periods (right). **(E)** Left: illustrations of rearing and rotating behavior. Right: group mean rates of rearing and rotating during the trial period.

**Figure 7.**
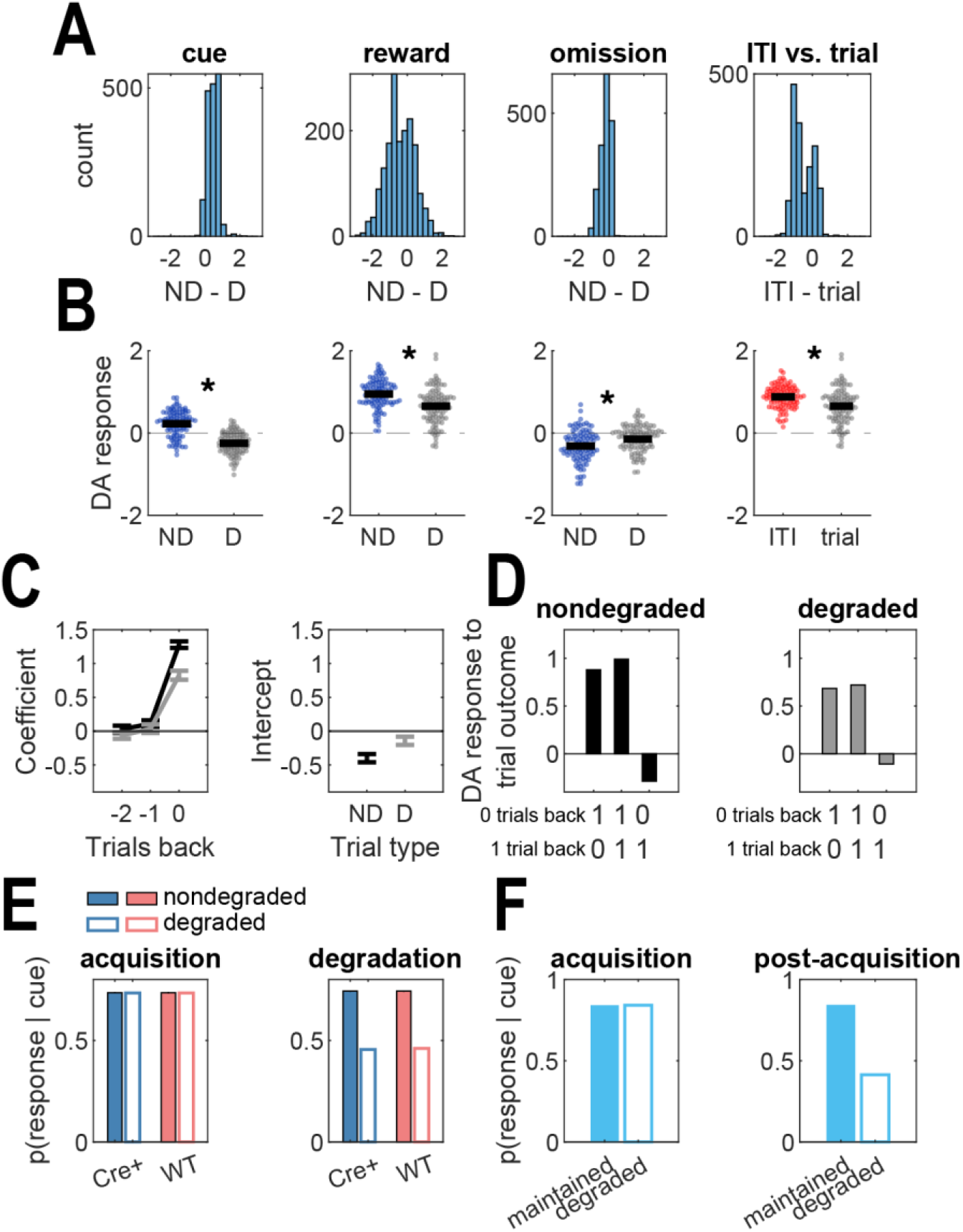
ANCCR simulations. **(A)** Distributions of simulated DA responses to trial events across all simulated parameter combinations. ND = non-degraded, D = degraded, ITI = degraded reward presented during the ITI, trial = degraded reward presented at the end of a trial. **(B)** DA responses to trial events from the best-fitting parameter combination. Means are taken from the final 50 trials of contingency degradation. *Separate paired t-test *p*’s < .0009 **(C)** Coefficients and intercepts when regressing the DA response to trial outcome on trial outcome history. Means are taken from the final 50 trials of degradation. **(D)** Predicted DA response at the time of the outcome on trial *n* as a function of the outcome on trial *n* and trial *n*-1. 1 indicates reward and 0 indicates omission. Means are taken from the final 50 trials of degradation. **(E)** Simulated behavioral response to the degraded cue when DA response to non-contingent rewards is inhibited (Cre+) or not (WT). **(F)** Behavioral simulation results when a DA response follows a 7 sec cue (acquisition) and is then also delivered noncontingently during the ITI (post-acquisition, degraded) or not (maintained).

TH-Cre+ rats expressing ChR2 acquired a modest level of conditioned locomotion during Pavlovian acquisition, but not enough to render a group x session interaction (**Figure 6C**; *F*’s(1,29) < 3.50, *p*’s > .073). This modest level of responding was ideal for achieving a sub-asymptotic level of behavioral responding to avoid overtraining. During post-acquisition, rats were split into groups that were either maintained on the same conditioning protocol or switched to contingency degradation. During contingency degradation, cues were still followed by laser stimulation, but ITI’s were filled with random deliveries of stimulation. The mean interval between stimulations during the ITI was designed to match the mean interval between cues and stimulation during trials.

Surprisingly, the degree of conditioned locomotion, measured as head velocity, did not differ between maintained and degraded groups during post-acquisition (**Figure 6C**). Repeated measures ANOVA’s detected significant effects of group (*F*’s(1,26) > 4.20, *p*’s < .027), but no main effects of session or group x session interactions. Post-hoc contrasts on the baseline-subtracted data revealed that the two TH-Cre+ groups showed greater conditioned locomotion than the TH-Cre-group during the final conditioning session (*p* < .05), but the two TH-Cre+ groups did not differ from each other (*p* > .05).

Locomotion was measured by tracking the velocity of the rats’ heads. There were other body parts that were tracked, and the angle, distance, and velocity between all pairs of points were computed for every frame. This created 135 different measurements per frame. The trial-averaged measures were compared between Cre groups for the final session of post-acquisition, and paired *t*-tests (α = .01) revealed a significant group difference only for three measures: the distances between the middle of the tail and the left ear, right ear, and middle of the head (**Figure 6D**). Each measure showed the same group patterns, so only the latter measurement was analyzed (trial – pretrial: *t*(20) = 2.88, *p* = .009). To understand the meaning of this group difference, each video from the final session of post-acquisition was hand-scored. Rats were timed for the duration of rearing and rotating during each trial (**Figure 6E**). An ANOVA revealed a main effect of behavior (*F*(1,20) = 5.44, *p* = .030) and a group x behavior interaction (*F*(1,20) = 6.07, *p* = .023). Post-hoc contrasts showed that the group that underwent contingency degradation showed a higher rate of rearing than rotating (*p* < .05), while the control group did not show a difference between the rate of the two behaviors (*p* > .05). Together, these results show that directly pairing a cue with VTA DA activity will elicit conditioned rotation only when the VTA DA activity is contingent on the cue.

### The effects of contingency degradation on DA signals and behavior are better explained by ANCCR (Adjusted Net Contingency for Causal Relations) than TDRL

It is difficult to explain many of our empirical observations during contingency degradation from the point of view of TDRL (**Supplemental Figure 9**). We therefore simulated a recently published reinforcement learning model called ANCCR (Adjusted Net Contingency for Causal Relations), which has successfully accounted for a range of phenomena related to Pavlovian conditioning and NAc dopamine release (Jeong et al., 2022). The model builds representations of contingency between pairs of events by learning how often a given event predicts all other events (successor representation), and how often a given event is preceded by all other events (predecessor representation). These representations are combined to yield the net contingency, and the dopamine response to an event is hypothesized to scale with the sum of all its net contingencies, plus the intrinsic meaningfulness of the event.

Model simulations reproduced the mean photometry transients aligned to trial events during contingency degradation (**Figure 7B**). The same model parameters also generated a pattern of regression coefficients that were similar to the empirical coefficients (**Figures 2C and 3C**), although there was one difference in that the non-degraded coefficient one trial back failed to reach a negative value (**Figure 7C**). The model also reproduced the results of the optogenetic experiments. The simulated behavioral response to the degraded cue was not affected by inhibiting the DA response during noncontingent rewards during the ITI (**Figure 7E**), as in the experiment in which VTA DA neurons were inhibited using eNpHR (**Figure 5**). And, like the experiment in which VTA DA neurons were stimulated using ChR2 (**Figure 6**), cue-evoked responding was attenuated when noncontingent DA responses were simulated during the ITI (**Figure 7F**). These results provide support for the ANCCR model in which DA signals whether an event causes another meaningful event (Jeong et al., 2022; see *Methods* for computational algorithm).

## Discussion

We provide experimental evidence showing that DA neuron activity in the midbrain and DA release in the ventral striatum are sensitive to changes in the contingency between specific cue-reward associations. We also demonstrate that non-contingent dopamine neuron activity is sufficient, but not necessary, to block a conditioned response. These findings pose a significant challenge to contiguity-based accounts of dopamine and learning, such as TDRL.

Delivering rewards during the ITI attenuated the Pavlovian conditioned response without inducing specific satiety or context overshadowing. It has long been known that Pavlovian learning depends on the contingency between cue and outcome, whether the outcome is appetitive (Delamater, 1995; Ostlund & Balleine, 2007) or aversive (Rescorla, 1968). During contingency degradation, the value of the cue remained unchanged because it was followed by the same reward at the same interval and the same probability as prior to the contingency manipulation. It is therefore difficult to explain the observed behavioral effect from the point of view of TDRL.

If phasic DA encodes a TDRL prediction error, then a strict prediction is that there should be no difference between the DA responses to events during degraded and non-degraded trials (**Supplemental Figure 9A**). Instead, VTA DA activity and NAc DA release showed weaker cue- and reward-evoked responses when the cue-reward contingency was degraded. Similarly, regressing the DA response to reward as a function of trial outcome history should yield coefficients that are similar between the two trial types according to TDRL (**Supplemental Figure 9B**). We found that photometry transients at the time of reward were sensitive to trial outcome history in a manner consistent with prediction error encoding only during nondegraded trials, and this sensitivity was notably absent during degraded trials (**Figures 2C, 3C**). TDRL also predicts that inhibiting DA during the ITI should have no effect on conditioned responding (**Supplemental Figure 9D**). Indeed, we found that inhibiting DA activity and release rendered conditioned responding unaffected (**Figure 5**), but this is also consistent with the predictions of ANCCR (**Figure 7E**). However, unlike TDRL, ANCCR can explain the outcome-selective effect of contingency degradation (**Figure 7E**). TDRL also predicts that stimulating DA during the ITI should slightly amplify conditioned responding (**Supplemental Figure 9E**), while ANCCR predicts an attenuation of responding (**Figure 7F**). The empirical data are consistent with the latter prediction (**Figure 6F**). As stated above, DA signaling during noncontingent rewards was not necessary for rats to learn outcome-selective contingency degradation. ANCCR has two possible explanations for this.

First, if we assume that optogenetic inhibition reduced the DA response to the baseline level, then inhibiting DA signaling during noncontingent rewards will not affect learning because food reward is innately meaningful regardless of its dopamine response, and thus noncontingent rewards will continue to be used to update net contingencies (see *Methods* for computational algorithm). This, however, makes it difficult to explain why we also observed that inhibiting DA release during reward blunted acquisition of an instrumental action (**Figure 5G**). However, if we assume that optogenetic inhibition reduced the DA response below baseline, then ANCCR can explain both the attenuation of instrumental learning during reward-paired DA inhibition and the null effect of DA inhibition during noncontingent rewards. This is because ANCCR decreases reward value when the DA response to reward is negative, since this signals that the reward is “overcaused” in the animal’s model of the world (Jeong et al., 2022). During instrumental learning, a consistently negative DA response to reward will decrease its value and blunt acquisition. During outcome-selective Pavlovian contingency degradation, a negative DA response to noncontingent reward will multiply with the decreasing net contingency between reward and cue (see *Methods*), and may even facilitate the effect of contingency degradation.

In addition, noncontingent DA stimulation was sufficient for rats to decrease a component of the conditioned response to a cue was also followed by DA stimulation. ANCCR predicts that, when a cue is directly paired with DA stimulation, the stimulation becomes a meaningful causal target and thus acts like a natural reward to drive learning. Just like with natural reward, presenting noncontingent DA stimulations during the ITI will reduce the predecessor representation between cue and stimulation and negatively affect conditioned responding.

The present dataset is not the first to challenge TDRL as an explanation for DA function. Experiments in rats have shown that VTA DA neurons mimic a prediction error that accounts for expectations concerning stimulus identity, not just value (Keiflin et al., 2019; Sharpe et al., 2017; Takahashi et al., 2017). These findings led to the proposal that VTA DA neurons compute sensory prediction errors as part of the successor representation algorithm, which learns the frequencies with which a given state is followed by every other state (Gardner, Schoenbaum, & Gershman, 2018). This learning algorithm is similar to TDRL, but it learns stimulus-stimulus associations and reward identities. ANCCR takes this idea a few steps further by quantifying a way for agents to also learn how frequently a given state is *preceded* by every other state and accounting for the base rate of that state—known as the predecessor representation contingency. The successor and predecessor representation contingencies are then combined to yield the net contingency between every pair of states. Indeed, our simulations show that, while the successor representation contingencies between the two cue-reward pairs was similar on average, the predecessor representation contingency was high at the time of reward following the nondegraded cue and low during reward following the degraded cue (**Supplemental Figure 10**).

These proposed underlying representations also help to explain why the DA response to reward following the degraded cue is smaller than to reward following the nondegraded cue. The ANCRR model considers DA to scale with the sum of net contingencies between a given event and all meaningful causal targets. During outcome-selective contingency degradation, both reward types are innately meaningful, and hence, the DA response to each reward type will depend on its contingency with respect to all other events. The predecessor representation contingencies between degraded rewards (i.e. the reward type delivered during the ITI) and all other events are negative (**Supplemental Figure 9**) because its base rate is high, and the predecessor representation contingency is negatively affected by a high base rate (see *Methods* for computational algorithm).

Two other studies have recorded neural activity during Pavlovian contingency degradation. One study of electrophysiological recordings in monkeys experiencing cues with high reward probabilities with and without ITI rewards found that the cue-evoked excitatory response of amygdala neurons was suppressed during contingency degradation, similar to the present work, but none of these neurons responded to reward delivery (Bermudez & Schultz, 2010). Another study of dLight recordings in the NAc in mice before and after Pavlovian contingency degradation found that the cue-evoked DA response was suppressed, as it was in our rats (Jeong et al., 2022). Of note, neither of these experiments used outcome-selective contingency degradation.

In summary, TDRL was insufficient to fully explain behavior and DA dynamics when contingency was manipulated. Instead, an alternative model based on estimates of causality explained almost every major aspect of the data set. The experiments in this report made use of contingency degradation following Pavlovian acquisition, but to test the robustness of the model it would also be useful to run the experiments in the reverse order, where contingency degradation comes first. It would also be useful to understand DA dynamics during contingency learning in an instrumental setting, for which there is evidence pointing to the importance of prefrontal cortical DA (Lex & Haber, 2010; Naneix et al., 2009). The ANCCR model, however, has not yet been formally extended to instrumental conditioning. Further developments in the model are therefore needed. The present work adds to a growing complement of studies expanding our understanding of dopamine’s role in learning (Burke et al., 2023; Coddington et al., 2023; Keiflin et al., 2019; Saunders et al., 2018; Seitz et al., 2022; Sharpe et al., 2017; Takahashi et al., 2023), with an emerging interpretation being that dopamine receives information about and feeds into not only prospective cognitive maps, but retrospective maps too.

## Acknowledgements

We thank Céline Drieu for helpful discussions on photometry analysis, Andy Dong for help with video scoring and constructing optic fibers, and Cecelia Shuai for help with running a subset of rats. We thank Gil Costa and Antonis Asiminas for making rat vector illustrations available on SciDraw (scidraw.io). This work was supported by National Institutes of Health grants F32 DA054767 (E.G.) and R01 DA035943 (P.H.J.).

## Materials and Methods

### Rats and surgeries

Thirteen TH-Cre rats were used for fiber photometry recordings in the VTA (6 females, 7 males). Rats underwent surgery 4-5 weeks prior to the beginning of the behavioral experiment. During surgery, a virus (1 μ1 of AAVDJ-EF1a-DIO-GCaMP6f diluted to 5 x 10^12^ particles/ml in PBS) was injected at the following coordinate relative to bregma: AP - 5.8, ML +0.7, DV - 8. An optic fiber (Doric Lenses) attached to an adapter was then lowered to 0.2 mm above the virus coordinate.

Fourteen Long Evans rats were used for fiber photometry in the NAc (6 females, 8 males). Rats underwent surgery 4-5 weeks prior to the beginning of the behavioral experiment. During surgery, a virus (1 μ1 of AAV5-hSyn-dLight1.2 diluted to 4.6 x 10^12^ particles/ml in PBS) was injected at the following coordinate relative to bregma: AP +1.75, ML +1.7, DV - 7. An optic fiber (Doric Lenses) attached to an adapter was then lowered to 0.2 mm above the virus coordinate.

Eight Long Evans rats were used for the context switch and food preference experiments (4 males, 4 females). These rats did not undergo surgery.

Nineteen Long Evans rats (10 TH-Cre+, 9 TH-Cre-) were used for the optogenetic inhibition experiment in the VTA (9 females, 10 males). Rats underwent virus injection and fiber implant surgeries 4 weeks prior to the beginning of the behavioral experiment. During surgery, a virus (AAV5-Ef1α-DIO-eNPHR3.0-eYFP diluted to 4 x 10^12^ particles/ml in PBS) was injected bilaterally in the VTA at the following coordinates relative to bregma: 1 μ1 at AP - 5.8, ML +/- 0.7, DV - 8.4; 0.5 μ1 at AP - 5.8, ML +/- 0.7, DV - 7.4. Optic fibers attached to ferrules were then lowered bilaterally to the VTA at the following coordinates at a 15 degree angle: AP - 5.8, ML +/- 2.71, DV - 7.76.

Seven TH-Cre+ rats were used for the optogenetic inhibition experiment in the NAc (3 females, 4 males). Rats underwent virus injection surgeries 11 weeks prior to the beginning of the behavioral experiment. During surgery, a virus (AAV1-hSyn1-SIO-eOPN3-mScarlet-WPRE diluted to 5 x 10^12^ particles/ml in PBS) was injected bilaterally in the VTA at the following coordinates relative to bregma: 1 μ1 at AP - 6.2, ML +/- 0.7, DV - 8.4; 0.5 μ1 at AP - 6.2, ML +/- 0.7, DV - 7.4; 1 μ1 at AP - 5.4, ML +/- 0.7, DV - 8.4; 0.5 μ1 at AP - 5.4, ML +/- 0.7, DV - 7.4. Eight weeks later, optic fibers attached to ferrules were then lowered bilaterally to the NAc at the following coordinates at a 10 degree angle: AP +1.75, ML +/- 2.96, DV - 6.7.

Twenty-nine rats were used for optogenetic stimulation in the VTA (17 females, 12 males). Rats underwent surgery 4-5 weeks prior to the beginning of the behavioral experiment. During surgery, a virus (AAV5-Ef1a-DIO-ChR2-eYFP diluted to 4.2 x 10^12^ particles/ml in PBS) was injected at the following coordinates relative to bregma: 1 μ1 at AP - 6.2, ML +0.7, DV - 8.4; 0.5 μ1 at AP - 6.2, ML +0.7, DV - 7.4; 1 μ1 at AP - 5.4, ML +0.7, DV - 8.4; 0.5 μ1 at AP - 5.4, ML +0.7, DV - 7.4. An optic fiber attached to a ferrule was then lowered to the following coordinate: AP - 5.8, ML +0.7, DV - 7.5.

All implants were secured to the skull with dental acrylic applied around skull screws, the base of the ferrule, and, in rats undergoing photometry recordings, the adapter.

### Pavlovian conditioning with food rewards

Pavlovian conditioning experiments with food rewards were conducted in plexiglass chambers with grid floors surrounded by a sound-attenuating cubicle (Med-Associates). Chambers were equipped with a food reward port that contained an infrared beam. Each time the beam was broken, a port entry was recorded.

Rats were food deprived to 85% of their *ad libitum* weight. Experiments began with two sessions of port training during which one type of food reward (45 mg grain pellet or 0.1 ml 20% sucrose) was delivered randomly into a food port every 60 seconds on average. Pellet deliveries were associated with the sound of the dispenser and a clinking of the pellet into the port, and sucrose deliveries were associated with the sound of the syringe pump and two clicks separated by 0.2 seconds. Each port training session contained different food rewards, but the same port was used for both reward types.

During the acquisition phase of Pavlovian conditioning, each food reward was preceded by a unique auditory cue (pure tone or white noise). When an auditory cue was presented, it lasted for 20 seconds and was followed by 0.5 probability of its associated reward. The ITI was drawn from an exponential distribution with a mean of 4 minutes and a range of 21 seconds and 14 minutes. A session lasted 70 minutes, and within a session each trial type was presented 8 times in random order. Although reward was probabilistic, rats were guaranteed to experience 4 rewarded trials and 4 non-rewarded trials of each type. Pavlovian acquisition lasted for 8 sessions.

During contingency degradation, the task contingencies remained the same except one type of food reward was delivered during the ITI non-contingently. Specifically, reward was delivered during the ITI every 20 seconds with 0.5 probability except when the upcoming trial was less than 20 seconds away. This was to avoid non-contingent rewards being delivered during the 20 second pre-cue baseline. Contingency degradation lasted for 8 sessions. Cue-reward assignments, non-contingent reward identity, and sex were counterbalanced.

Following contingency degradation, one session of reacquisition was given during which non-contingent rewards were no longer delivered during the ITI.

### Context switch

Rats were put through the same series of Pavlovian conditioning and contingency degradation sessions as described above. Following the last session of degradation, two extinction sessions were conducted during which 4 non-rewarded tone and noise trials were presented in random order and at random times (exponentially distributed ITI with a mean of 4 mins). One session was conducted in the normal context and the other in a modified context, with an additional session of contingency degradation separating the two tests. The modified context had a floor made from plastic with raised points in a honeycomb pattern, stripe patterned walls, and lemon scent. The normal context had grid floors without the stripped patterns or any special scent. Cue-reward assignments, non-contingent reward identity, and sex were counterbalanced. Order of testing was counterbalanced with cue-reward assignments and non-contingent reward identity.

### Food preference tests

Following the context tests, two food preference tests were conducted during which rats were free to eat and drink the same pellet and sucrose rewards used in the Pavlovian conditioning experiment. The first test was conducted immediately after an additional session of contingency degradation. The second test was conducted the following day without any prior experimental session. Tests lasted for 30 minutes. The weights of the pellets and sucrose were recorded before and after the tests.

### Fiber photometry

A fluorescence mini-cube (Doric Lenses) transmitted light streams from a 465-nm LED sinusoidally modulated at 330 Hz that passed through a GFP excitation filter, and a 405-nm LED modulated at 120 Hz. Both 465 and 405 nm streams were bandpass filtered. LED power was set at 28 µW for the 405 stream and 68 µW for the 465 stream. GCaMP6f fluorescence from neurons below the fiber tip in the brain was transmitted via this same low-autofluorescence fiber cable (400 nm, 0.52 NA) back to the mini-cube, where it was passed through a GFP emission filter, amplified, and focused onto a high sensitivity photoreceiver (Doric Lenses). A real-time signal processor (RZ5P, Tucker-Davis Technologies) running Synapse software modulated the output of each LED and recorded photometry signals, which were sampled from the photodetector at 6 kHz. The signals generated by the two LEDs were demodulated and decimated to 1020 Hz for recording to disk.

For analysis, signals were downsampled to 102 Hz, and the 465 nm signal was normalized to the 405 nm signal by computing ΔF/F. Specifically, the best-fitting line relating the 465 nm signal to the 405 nm signal was estimated, and the 405 nm signal was then transformed by multiplying by the regression slope and adding on the y-intercept. This puts the 405 nm data in the range of the 465 nm data. ΔF/F was computed as (465-nm – transformed 405-nm) / (transformed 405-nm).

### Optogenetic inhibition of VTA DA neurons during Pavlovian contingency degradation

Rats with optic fibers targeting the VTA were put through the identical experiment described above (see *Pavlovian conditioning with food rewards*). During contingency degradation rats received green laser stimulation delivering 4 seconds of constant 532 nm light (12-15 mW) at the onset of every noncontingent reward delivered during the ITI. Rats were tethered to a bilateral patch cord during every session of the experiment.

### Optogenetic inhibition of DA terminals in NAc during Pavlovian contingency degradation

Rats with optic fibers targeting eOPN3-expressing VTA axons in the NAc were put through a similar behavioral experiment described above (see *Pavlovian conditioning with food rewards*). During contingency degradation rats received green laser stimulation delivering 1 second of constant 532 nm light (10 mW) at the onset of the first reward delivered during the ITI. The effect of eOPN3 activation lasts for minutes (Mahn et al., 2021), and this creates a tradeoff between minimizing the eOPN3 activation that continues into the next trial and the number of rewards during the ITI that are given in the absence of eOPN3 activation. To strike a reasonable balance between this tradeoff, the range of the ITI’s was changed to 2-6.5 minutes. The mean ITI was maintained at 4 minutes. In addition, no rewards were delivered during the ITI when the upcoming trial was less than 60 seconds away. Rats were tethered to a bilateral patch cord during every session of the experiment.

### Optogenetic inhibition of DA terminals in NAc during instrumental acquisition

Rats that underwent inhibition of DA terminals in the NAc during Pavlovian contingency degradation were next trained to press two levers on fixed-ratio 1 schedules in separate sessions. Two 30-minute sessions were run each day for a total of three days during which one lever was presented and rats were free to press the lever for one grain pellet reward at any time. One lever was assigned as the laser lever and the other was assigned as the control lever. When the laser lever was pressed, a pellet reward was triggered along with 1 second of constant 532 nm light (10 mW). Laser was withheld if the previous laser onset occurred within the last sixty seconds. The control lever session did not include any laser stimulation, although rats were still tethered to the patch cord. The order of testing was balanced across rats and alternated each day.

### Pavlovian conditioning with optogenetic stimulation

Rats with fibers targeting ChR2-expressing neurons in the VTA were tethered to a patch cord connected to a laser via a commutator. Lasers delivered 473 nm blue light (12 mW).

Conditioning was divided into two phases: acquisition and post-acquisition. During acquisition, a compound cue (panel light and pure tone) was presented for 7 seconds followed by 1 second laser stimulation (20 5-ms pulses at 20 Hz). ITI’s were drawn from an exponential distribution with a mean of 200 seconds and a range of 15 to 615 seconds. Sessions lasted for 86 minutes during which 25 trials were presented. Rats were split into two groups during acquisition: Cre-(4 females, 3 males) and Cre+ (14 females, 10 males).

During post-acquisition, some rats were maintained on the same conditioning protocol (Cre+: 7 females, 3 males; Cre-: 1 female, 2 males) while others underwent contingency degradation (Cre+: 6 females, 6 males; Cre-: 3 females, 1 male). All experiment parameters remained identical except non-contingent 1 second laser stimulations were delivered during the ITI with a probability of 0.125 every second. Two Cre+ rats from the acquisition phase were dropped from post-acquisition analysis because one became disconnected from the patch cords during the last session and the other scratched its head to the point of bleeding prior to the start of the last session.

### Histology

At the end of all experiments, rats were perfused transcardially with 0.9% saline followed by 4% paraformaldehyde (PFA). Brains were removed and stored in PFA for 1 hour followed by 30% sucrose in PBS for 72 hours. Coronal sections 50 µm thick were cut using a cryostat, and sections were stored in PBS at 4° C. Brain sections were mounted on microscope slides, coverslipped with VECTASHIELD Antifade Mounting Medium with DAPI, and examined with a fluorescent microscope (Zeiss).

### Statistical analysis

Statistical tests on summary data included repeated measures ANOVA and *t*-tests with a significance threshold of 0.05 except where noted in the main text. Significant interactions were followed up with post-hoc contrasts using the recommendations of Rodger (1974).

All ΔF/F photometry traces were z-scored relative to a pre-trial 20 second baseline. When z-scoring non-contingent rewards during the ITI, the nearest pre-trial baseline was used. To summarize photometry responses to cues, the z-scored ΔF/F was averaged starting from the time of cue onset and ending 5 seconds later. To divide cue-evoked responses into phasic and tonic components, the phasic beginning and end periods were defined as the times after cue onset at which the lower bound of a 95% confidence interval passed above and dropped back down to 0, respectively (Jean-Richard-dit-Bressel, Clifford, & McNally, 2020). For the VTA GCaMP6F experiment, this interval was between 255 ms and 549 ms after cue onset. For the NAc dLight experiment, this interval was between 245 ms and 372 ms after cue onset. The tonic component of the cue-evoked responses was the mean of the photometry signal starting from the end of the phasic window and ending right before cue offset.

To summarize photometry responses to rewards, the z-scored ΔF/F was averaged starting from the time of reward onset and ending 10 or 5 seconds later for VTA GCaMP6f and NAc dLight, respectively.

Reward omission responses were divided into positive and negative phases. The positive phase was averaged starting from the moment after cue offset and ending when activity significantly dropped below baseline. The negative phase was averaged starting from the moment after cue offset when activity significantly dropped below baseline and ending when activity rose back up to baseline. To identify when the signal significantly dropped below baseline, the time at which the lower bound of a 95% confidence interval dropped below 0 was used (Jean-Richard-dit-Bressel, Clifford, & McNally, 2020). For the VTA GCaMp6f experiment, the averaging window for the negative phase was between 3.31 and 9.76 seconds post-trial. For the NAc dLight experiment, the window was between 0.66 and 2.29 seconds post-trial.

To quantify how local reward history influenced conditioned responding and photometry responses to cues and outcomes, we used multiple regression. The number of cued port entries on trial *t* was modeled as a linear combination of trial outcomes:

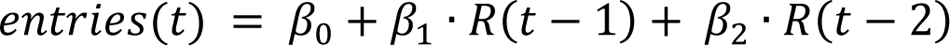

where *R* is trial outcome (1 if rewarded, 0 if omitted). The GCaMP and dLight responses to trial outcome and cue was modeled in a similar way:

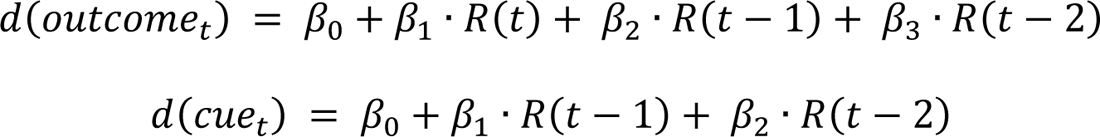

The trial outcome was averaged over the start and end points used to define the reward omission response window (see previous paragraph).

To predict conditioned behavior from photometry signals on a trial-by-trial basis, three multiple regression analyses used the mean z-scored ΔF/F response to cue and trial outcome, as well as trial number:

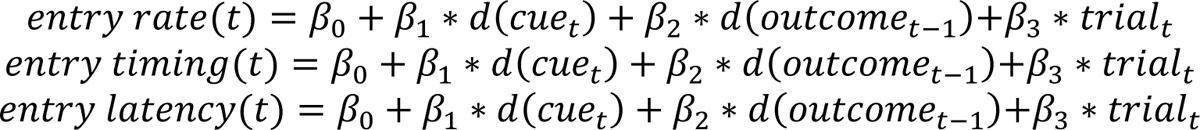

Entry rate was defined as total number of entries during the 20 second cue. Entry timing was defined as the area under the curve of the cumulative proportion of entries during the 20 second cue. Entry latency was defined as the time from cue onset to the first entry during the 20 second cue.

Behavioral data during conditioning with optogenetic stimulation was derived from body part location estimates using DeepLabCut (Mathis et al., 2018). Each video frame contained the estimated x-y positions of the following body parts and environmental cues: left ear, right ear, middle of the head between the ears, middle of the back, base of the tail, middle of the tail, four corners of the behavioral chamber, and the left panel light, which functioned as part of the compound cue. DeepLabCut also generated a confidence measure ranging from 0 to 1, and trials containing frames with confidence measures less than 0.95 were excluded from analyses. To derive the velocity of the rat, the distance between the middle of the ears and the panel light was measured in pixels for each frame, and the differences between the frame-by-frame distances were computed. Video scoring was performed by using a stopwatch to quantify the duration of rearing and rotating during each trial.

### ANCCR simulations

All simulations were performed in MATLAB with the help of functions available on the Namboodiri lab GitHub (https://github.com/namboodirilab/ANCCR). Simulated experimental events mimicked the same contingencies during acquisition and contingency degradation, except the number of trials was increased to 500 per cue in each phase. We first simulated ANNCR on the photometry experiments, using twenty iterations per parameter combination. There were six free parameters set to the following ranges: *T ratio* = 0.2-1.4, *α* = 0.01-0.3, *k* = 0.01-0.6, *w* = 0.1-0.7, *threshold* = 0.1-0.7, *α_R_* = 0.1-0.3. The winning parameter combination was determined by maximizing the correlation between the rankings of the following empirical and simulated means: ITI reward, degraded reward, nondegraded cue, and degraded cue. Once the winning parameter combination was identified, the model was simulated again for 100 iterations.

The winning set of parameters were then used to simulate behavioral responding during the optogenetic inhibition and stimulation experiments using 100 iterations per experiment.

ANCCR computations are described below. Event *i* is kept in memory across time steps according to an eligibility trace

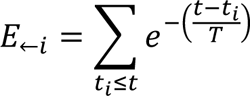

where *T* is the temporal decay parameter. *T* was always considered a fraction of the mean inter-reward interval during acquisition, and this fraction (*T ratio*) was a free parameter used to fit the model.

ANCCR involves computing the adjusted net contingency between all pairs of events and using that to generate DA and behavioral responses. The adjusted net contingency is derived from the net contingency, which is derived from the predecessor and successor representation contingencies. The predecessor representation between events *i* and *j* is updated at the time of *j* as

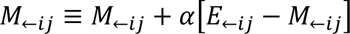

where *E*_←ij_ measures the eligibility trace of *i* are the current time of *j* and ≡ denotes an update operation. The predecessor representation quantifies how often the event *i* precedes the event *j*. The agent also keeps track of the baseline rate of all events according to where – represents random moments. We assumed it is updated every 0.2 s. The predecessor representation contingency (PRC) is then calculated as

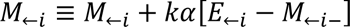

The successor representation contingency (SRC) is then calculated using Bayes’ rule and is updated as the time of *i*:

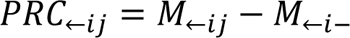

The predecessor and successor representation contingencies were then combined into a weighted sum to yield the net contingency (NC):

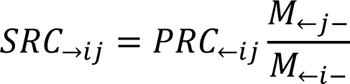

The adjusted net contingency (ANCCR) between *i* and *j* is then calculated to account for possible causes of *i*, like when it is consistently preceded by *k*:

Here, Δ measures recency of *k* with respect to *i* and is defined as R_ij_ is a causal weight that is updated according to

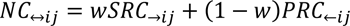

where δ_ij_ is a prediction error that is computed according to

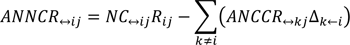

where R_jj_ is the externally signaled reward magnitude of *j* and *n_i_* is the total number of times event *i* has been experienced. The dopamine response to an event *i* is the sum of learned meaningfulness of stimlulus *i* (ANCCRs of *i* with respect to all meaningful causal targets *j*) and the innate meaningfulness (*b*_i_):

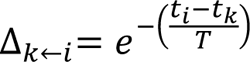

In the original proposal (Jeong et al., 2022), the dopamine response was defined as the learned meaningfulness of a stimulus, but it has been revised to account for both the learned and innate meaningfulness of a stimulus in a subsequent paper (refer to Berke et al., 2023 for the rationale behind this model update).

For *j* to be considered a meaningful causal target, the DA response must pass a threshold:

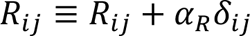

The variable *b*_j_ represent the innate meaningfulness of stimulus *j* and was always set to 0.5 for rewards and 0 for all other stimuli. To generate behavioral response probabilities, we applied a softmax function:

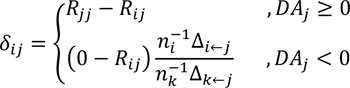

where *β* is the inverse temperature and was set to 5. *V* is the value of a cue and was defined as *NC*_↔*ij*_ R_*ij*_ − Cost. Cost was set to 0.3.

The set of parameters that best fit the photometry data was *T ratio* = 1, *α* = 0.2, *k* = 0.01, *w* = 0.4, *threshold* = 0.7, *α_R_* = 0.1.

### TDRL simulations

We simulated TDRL on the same experiments as ANCCR. Event *i* is kept in memory across time steps *t* according to an eligibility trace

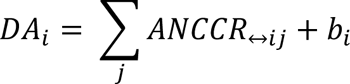

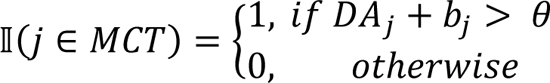

where *x*(*i*)_*t*_ is the activation of state *i* associated with event *x* at time *t*. The representation of events as discrete non-overlapping states is known as the complete serial compound (Schultz et al., 1997). The values of each state at time *t* are a weighted sum of their activations at time *t*:

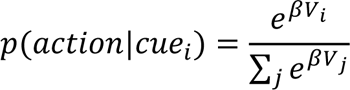

Weights are updated according to the size of the temporal difference prediction error, shown in brackets:

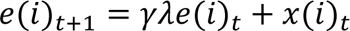

Under this model, the DA response to any given event is equal to the prediction error at the time of the event. Behavioral response probabilities were generated according to:

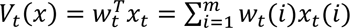

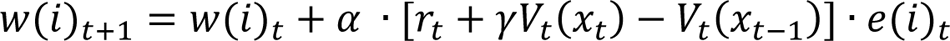

where *β* is the inverse temperature and was set to 5. *V* is the value of a cue as defined above (*V*_*t*_(*x*)). Cost was set to 0.3. γ was set to 0.85 and λ was set to 0.

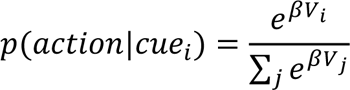

---

**Supplemental Figure 1.**
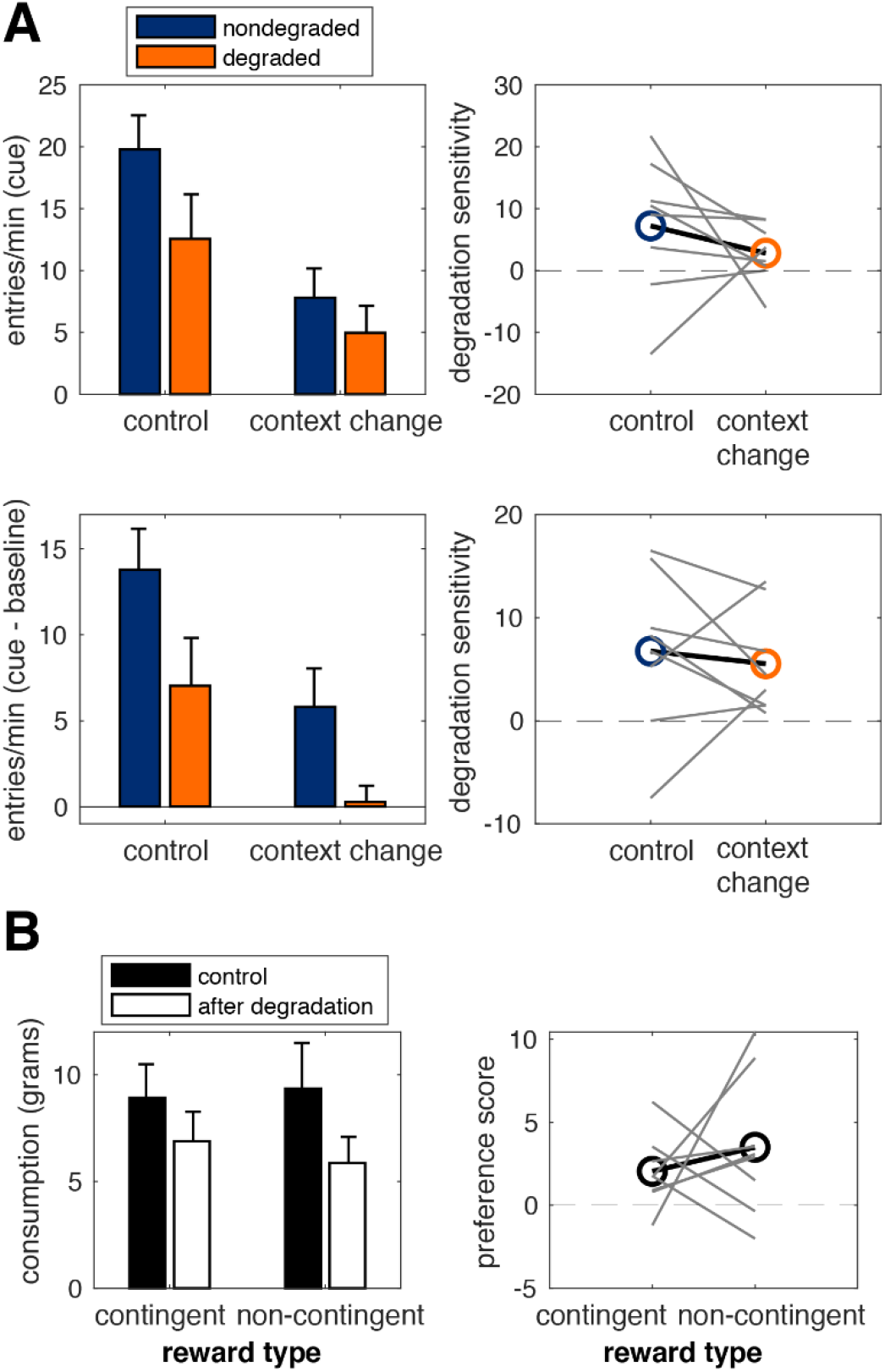
**(A)** Left: port entry rates during the cue (top) and with baseline subtracted (bottom) during extinction tests conducted in the original conditioning context (left bars) and the altered context (right bars). Right: degradation sensitivity scores taken as a difference between nondegraded and degraded entry rates. Main effect of test condition (*p*’s = .048), main effect of trial type (*p*’s = .042), no test x trial interaction (*p*’s > .364). **(B)** Left: consumption of rewards after a session of contingency degradation (white bars) versus no prior conditioning session (black bars). The non-contingent reward is the one given in the absence of the cue, while the contingent reward is the one given only at the end of a trial Right: preference scores calculated as the difference between consumption during control session and the session that occurs after contingency degradation. Main effect of test condition (*p* = .003), no main effect of reward type (*p* = .911), no test x reward interaction (*p* = .499). contingency degradation. Main effect of test condition (*p* = .003), no main effect of reward type (*p* = .911), no test x reward interaction (*p* = .499).

**Supplemental Figure 2.**
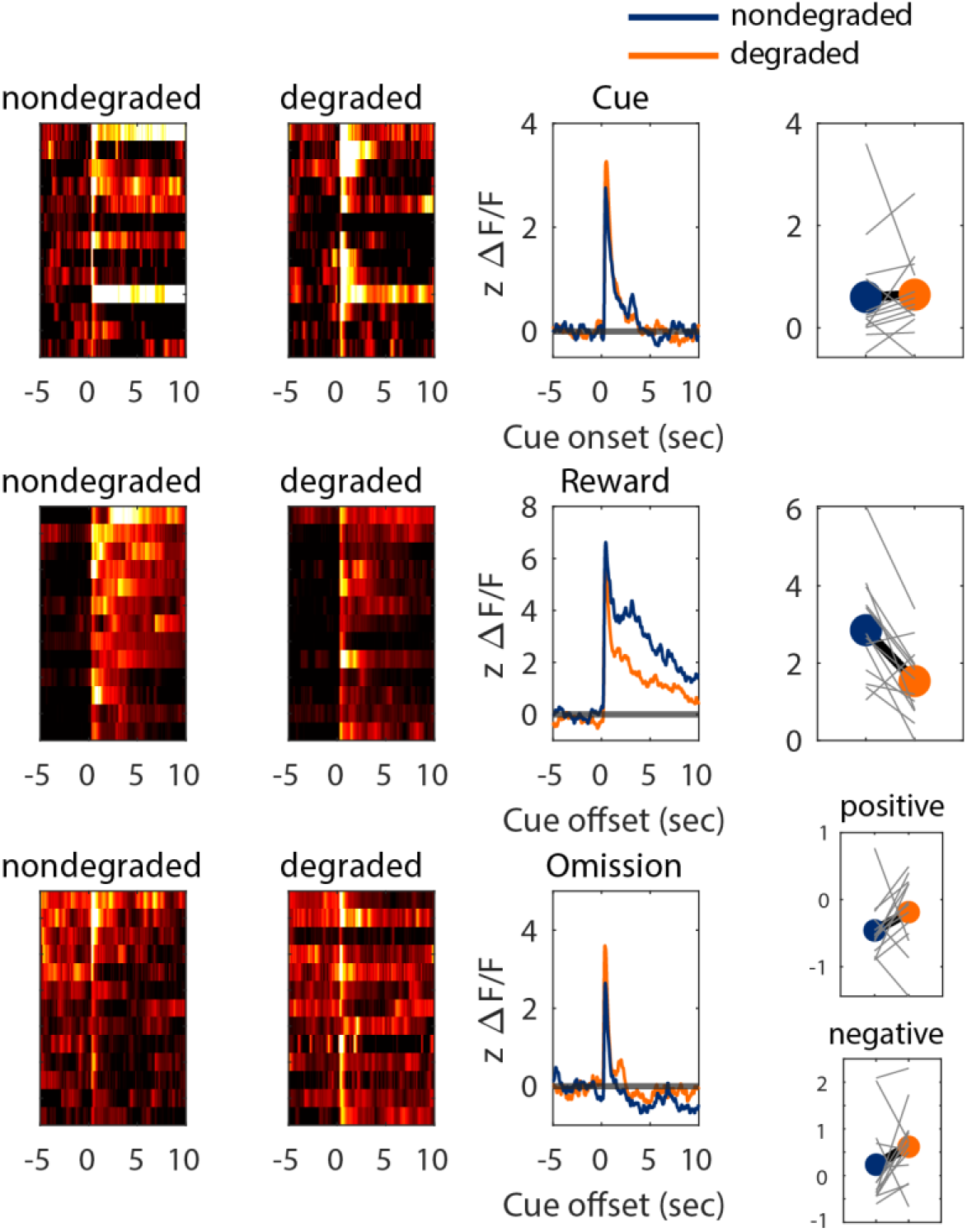
VTA GCaMP data from first session of contingency degradation. Left column: heat maps showing GCaMP response aligned to trial events separated by nondegraded and degraded trials. Each row shows the mean for each rat. Middle column: Mean photometry traces in response to trial events, shown separately for nondegraded (blue) and degraded (orange) trials. Right column: mean z-scored ΔF/F in response to trial events for nondegraded and degraded trials. Reward omission means were quantified separately for positive (top) and negative (bottom) phases of the signal. Grey lines show individual rats.

**Supplemental Figure 3.**
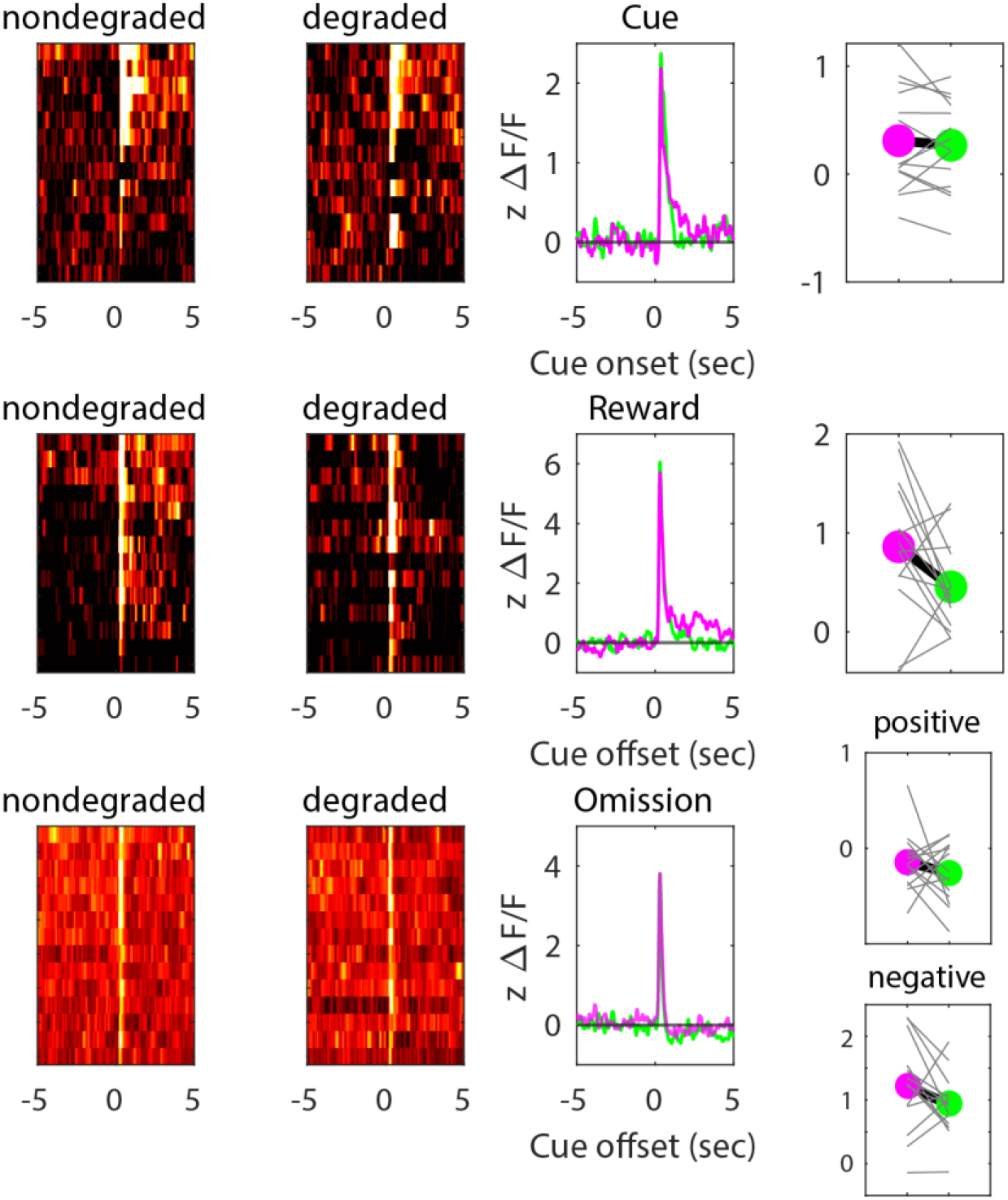
NAc dLight data from first session of contingency degradation. Left column: heat maps showing dLight response aligned to trial events separated by nondegraded and degraded trials. Each row shows the mean for each rat. Middle column: Mean photometry traces in response to trial events, shown separately for nondegraded (blue) and degraded (orange) trials. Right column: mean z-scored ΔF/F in response to trial events for nondegraded and degraded trials. Reward omission means were quantified separately for positive (top) and negative (bottom) phases of the signal. Grey lines show individual rats.

**Supplemental Figure 4.**
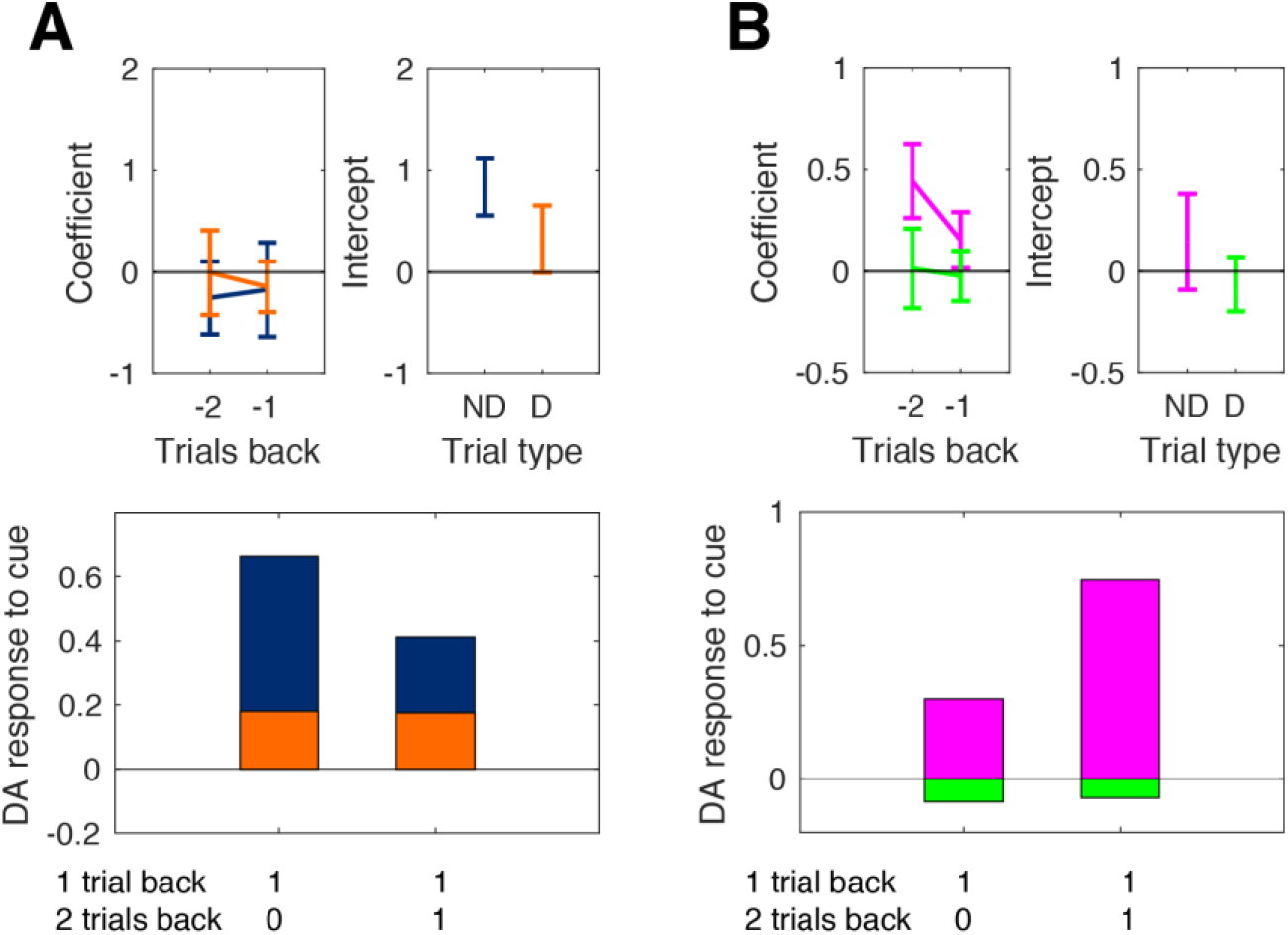
Cue-evoked photometry signals were regressed against trial outcome history. Top: mean regression coefficients (left) and intercepts (right) when the photometry response to the cue was regressed against trial outcome history, shown separately for nondegraded and degraded trials. Bottom: mean predicted photometry response at the time of the cue on trial *n* as a function to the identity of the outcome on trial *n*-1 and trial *n*-2. 1 indicates reward and 0 indicates omission. All data are taken from the last session of contingency degradation. (**A**) VTA GCaMP data. No main effect of cue (*F*(1,12) = 1.08, *p* = .318). No main effect of trials back (*F*(1,12) = .24, *p* = .630). No cue x trials back interaction (*F*(1,12) = .18, *p* = .678). **B** NAc dLight data. Main effect of cue (*F*(1,13) = 12.10, *p* = .004). Main effect of trials back (*F*(1,13) = 4.72, *p* = .049). No cue x trials back interaction (*F*(1,13) = 1.88, *p* = .193).

**Supplemental Figure 5.**
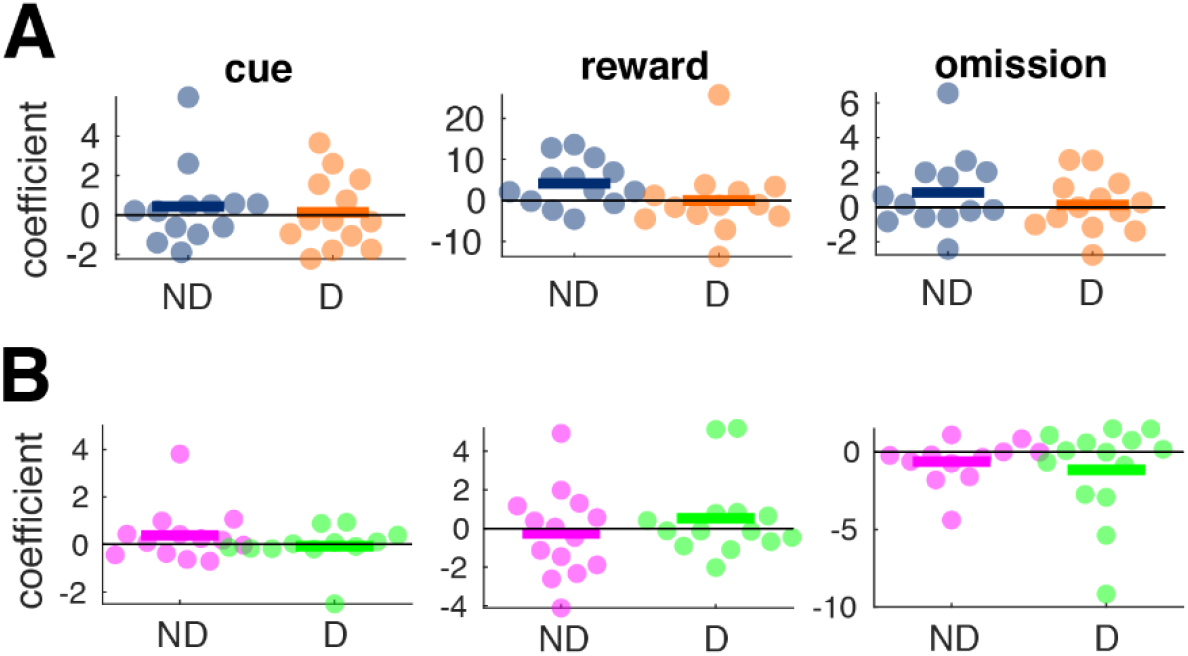
Photometry signals evoked by cue, reward, and reward omission were regressed against the time in session at which the event occurred. Each data point is an individual rat and horizontal bars are means. Only the negative portion of the photometry response to omission is shown. (**A**) VTA GCaMP data. (**B**) NAc dLight data. ND = non-degraded, D = degraded. All data are from the last session of contingency degradation.

**Supplemental Figure 6.**
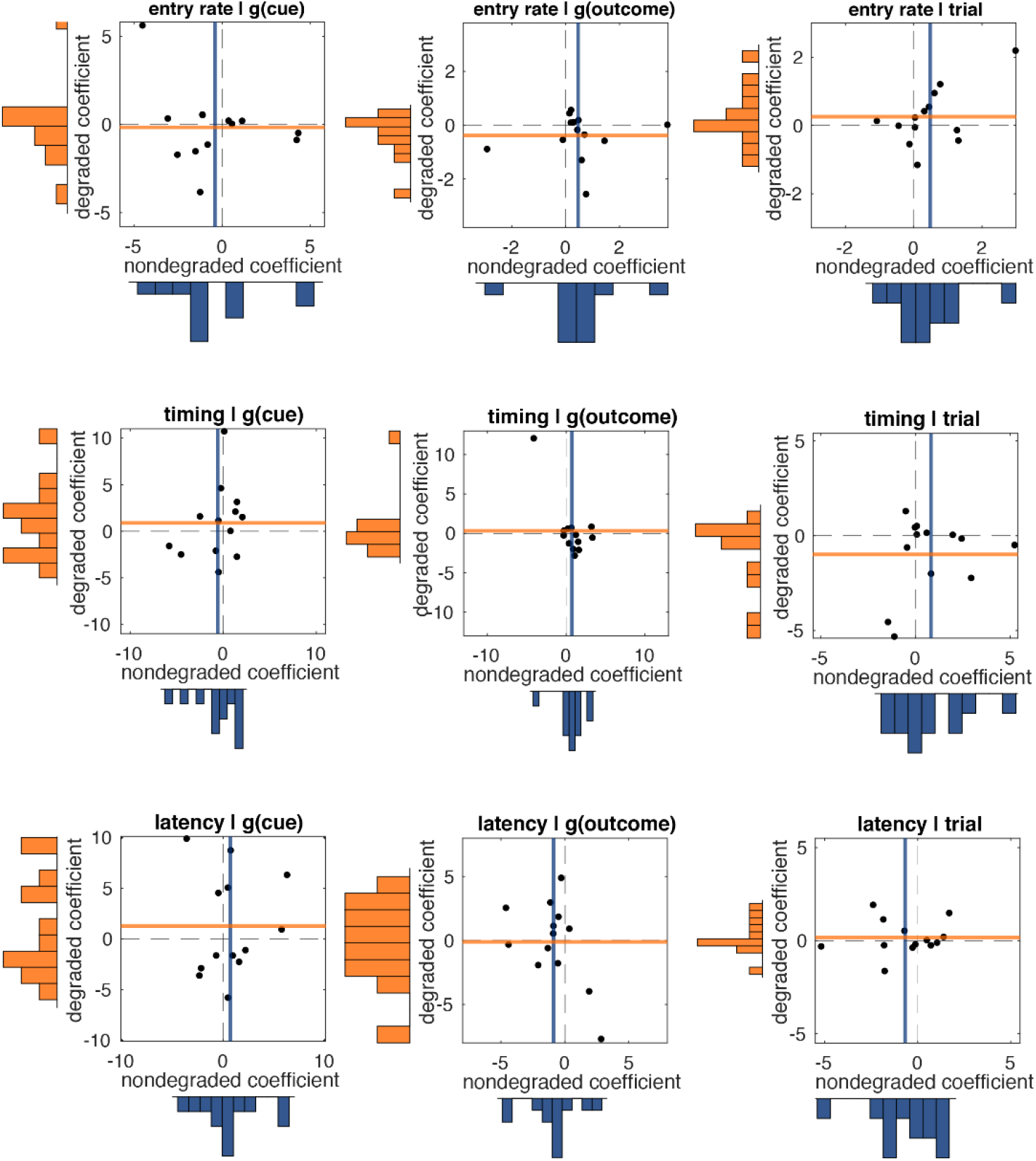
Distributions of coefficients in the VTA GCaMP experiment when the entry rate (top row), entry timing (middle row), and entry latency (bottom row) were regressed against the GCaMP response to the cue on the same trial (left column), the GCaMP response to the trial outcome on the previous trial (middle column), and trial number (right column). Each row represents the results from a single multiple regression analysis. All data are from the last session of contingency degradation.

**Supplemental Figure 7.**
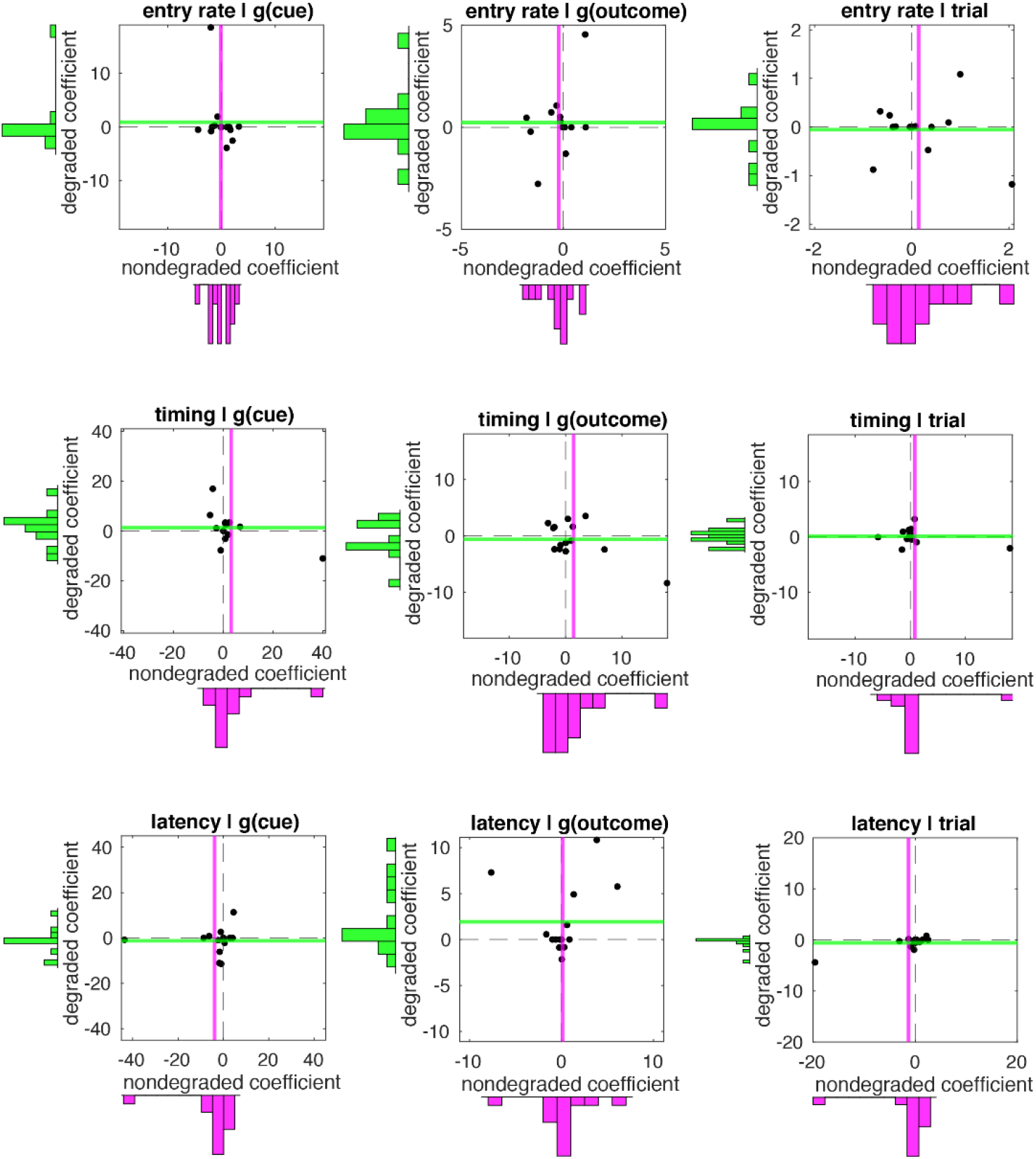
Distributions of coefficients in the NAc dLight experiment. The description of the figure is the same as Supplementary Figure 6.

**Supplemental Figure 8.**
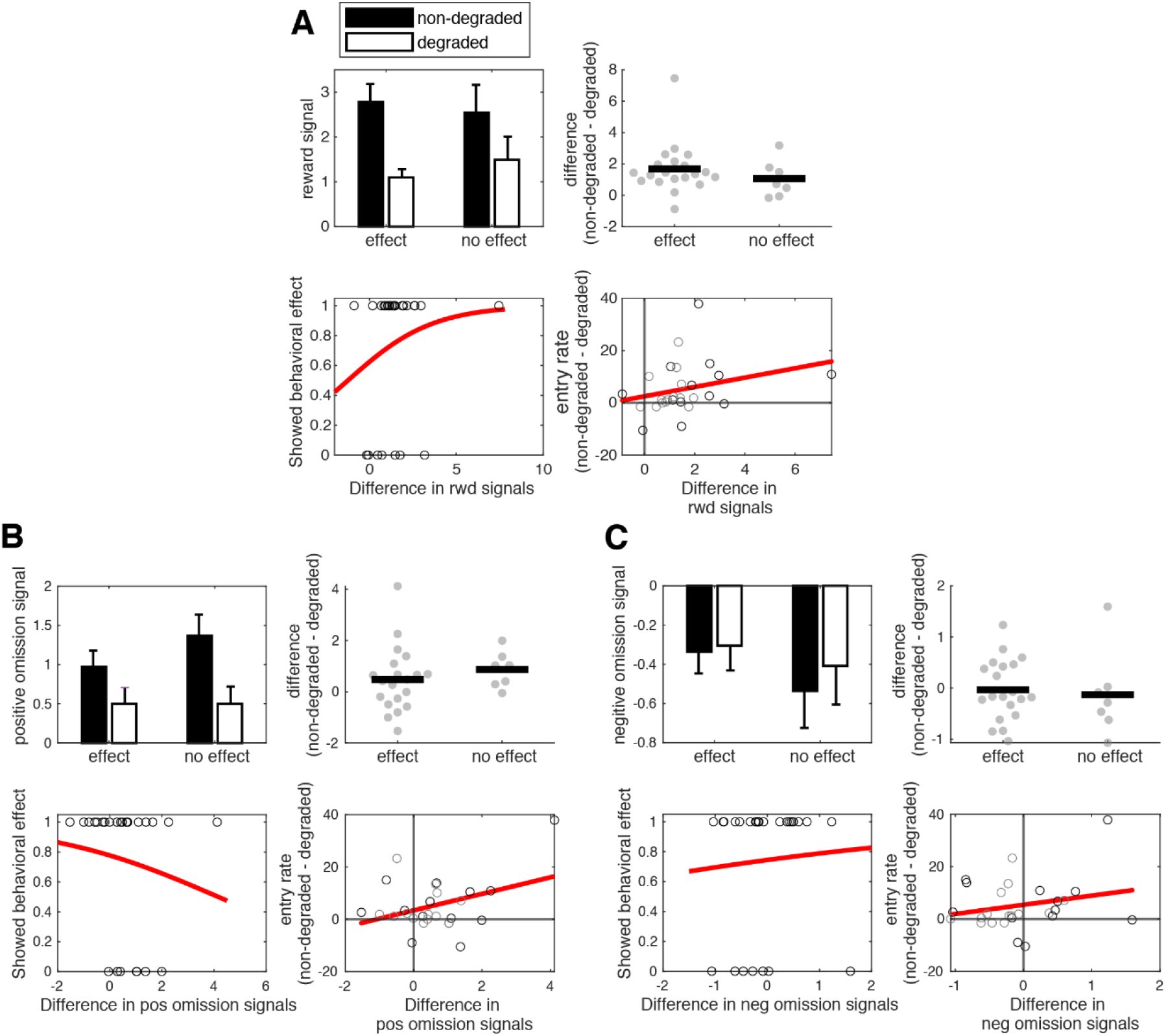
Rats from the VTA GCaMP and NAc dLight experiments were combined and then split into groups that showed an effect of contingency degradation and those that did not. The photometry responses to reward (**A**) and omission (**B and C**) at the end of nondegraded and degraded trials could not discriminate between groups. All data are from the last session of contingency degradation.

**Supplemental Figure 9.**
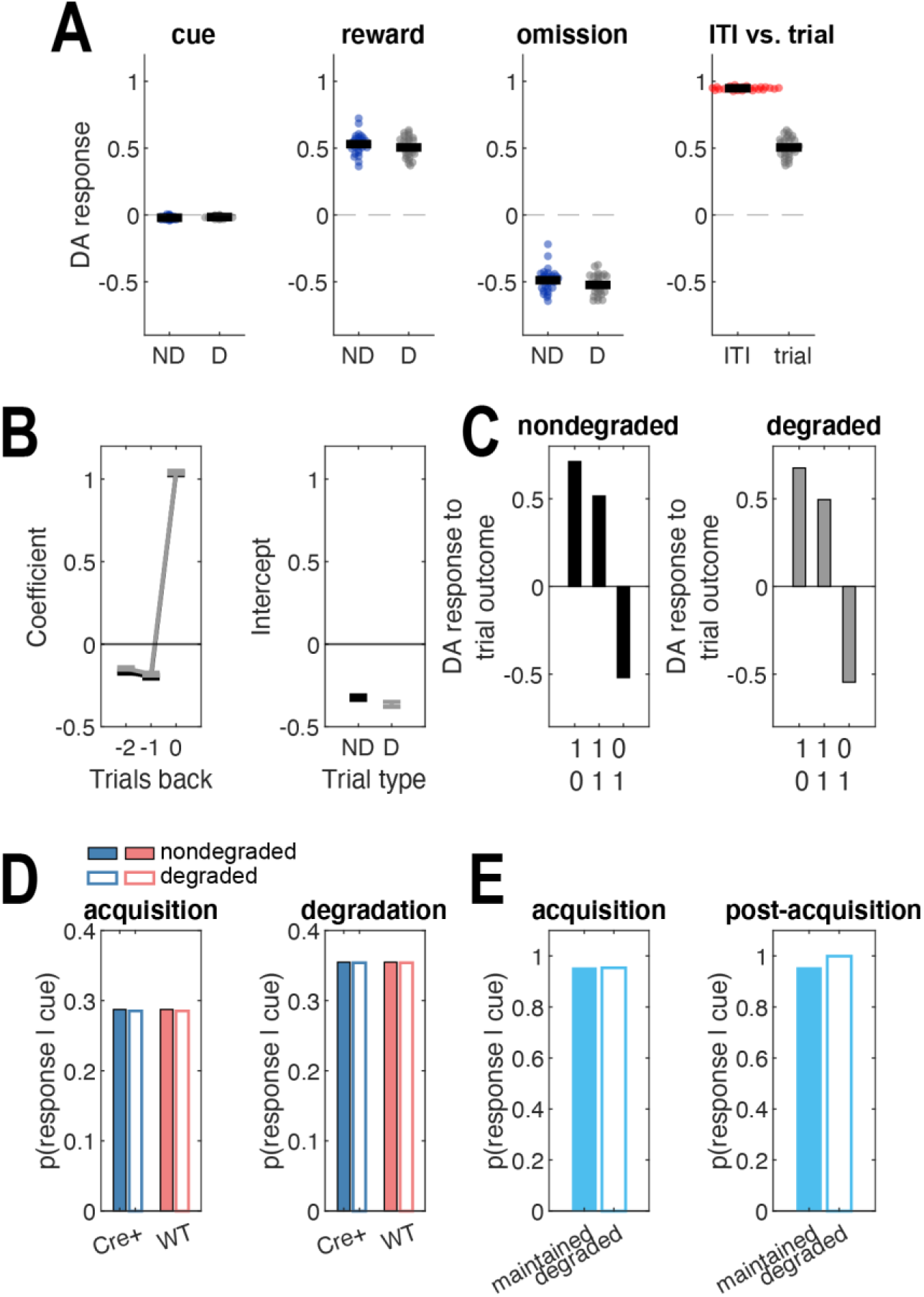
TDRL simulations. **(A)** DA responses to trial events. Means are taken from the final 50 trials of contingency degradation. **(B)** Coefficients and intercepts when regressing the DA response to trial outcome on trial outcome history. Means are taken from the final 50 trials of degradation. **(C)** Predicted DA response at the time of the outcome on trial *n* as a function of the outcome on trial *n* and trial *n*-1. 1 indicates reward and 0 indicates omission. Means are taken from the final 50 trials of degradation. **(D)** Simulated behavioral response to the degraded cue when DA response to non-contingent rewards is inhibited (Cre+) or not (WT). **(E)** Behavioral simulation results when a DA response follows a 7 sec cue (acquisition) and is then also delivered noncontingently during the ITI (post-acquisition, degraded) or not (maintained).

**Supplemental Figure 10.**
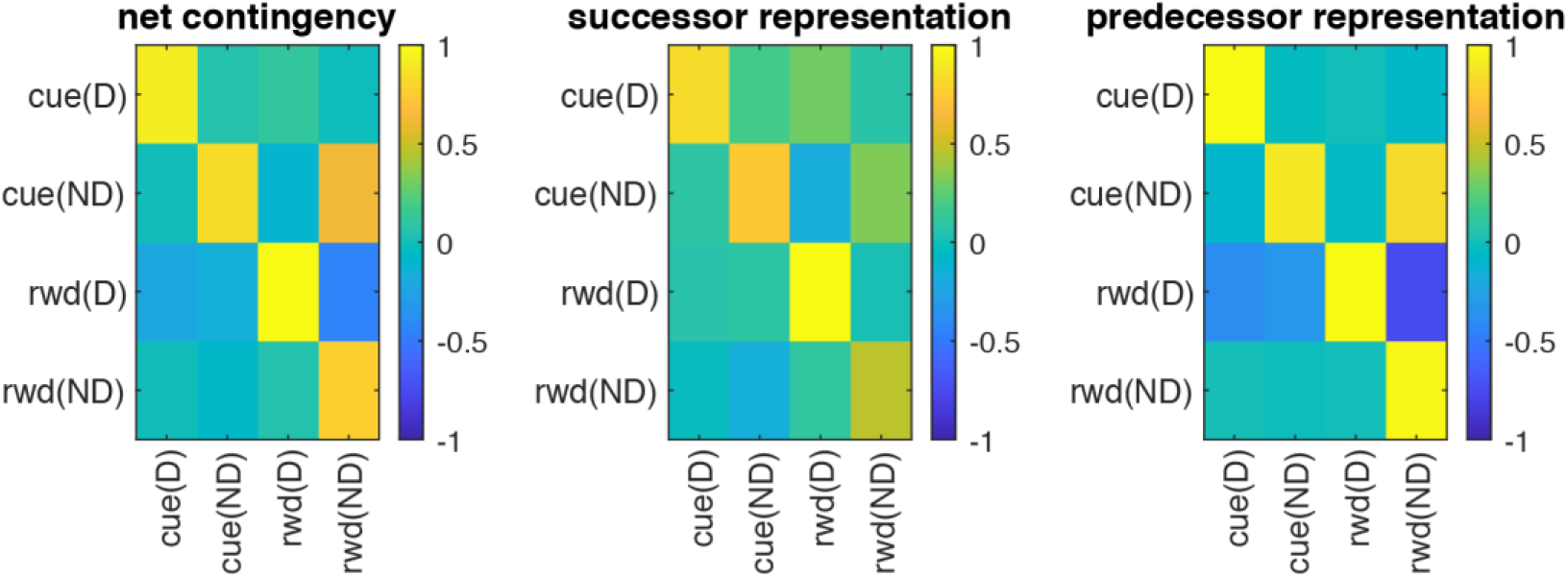
Results from ANCCR simulations showing the magnitudes of net contingency, successor representation contingencies, and predecessor representation contingencies between all events during outcome-selective contingency degradation. Data are averaged over the final 50 trials of contingency degradation. D = degraded, ND = non-degraded, rwd = reward. Under ANCCR, the DA response to an event is proportional to the sum of its net contingencies (left). The net contingency between a pair of events is a weighted sum of their successor and predecessor representation contingencies (middle and right).

